# Overlapping stimulons arising in response to divergent stresses in *Escherichia coli*

**DOI:** 10.1101/2021.12.17.473201

**Authors:** Huijing Wang, GW McElfresh, Nishantha Wijesuriya, Adam Podgorny, Andrew D. Hecht, J. Christian J. Ray

## Abstract

Cellular responses to stress can cause a similar change in some facets of fitness even if the stresses are different. Lactose as a sole carbon source for *Escherichia coli* is an established example: too little causes starvation while excessive lactose import causes toxicity as a side-effect. In an *E. coli* strain that is robust to osmotic and ionic differences in growth media, B REL606, the rate of antibiotic-tolerant persister formation is elevated in both starvation-inducing and toxicity-inducing concentrations of lactose in comparison to less stressful intermediate concentrations. Such similarities between starvation and toxification raise the question of how much the global stress response stimulon differs between them. We hypothesized that a common stress response is conserved between the two conditions, but that a previously shown threshold driving growth rate heterogeneity in a lactose-toxifying medium would reveal that the growing fraction of cells in that medium to be missing key stress responses that curb growth. To test this, we performed RNA-seq in three representative conditions for differential expression analysis. In comparison to nominally unstressed cultures, both stress conditions showed global shifts in gene expression, with informative similarities and differences. Functional analysis of pathways, gene ontology terms, and clusters of orthogonal groups revealed signatures of overflow metabolism, membrane component shifts, and altered cytosolic and periplasmic contents in toxified cultures. Starving cultures showed an increased tendency toward stringent response-like regulatory signatures. Along with other emerging evidence, our results show multiple possible pathways to stress responses, persistence, and possibly other phenotypes. These results suggest a set of overlapping responses that drives emergence of stress-tolerant phenotypes in diverse conditions.

## 1. Introduction

Fitness and survival of a single-celled species across diverse environments incur classic trade-offs between metabolic costs and improvement of survival probability: an effective response in one environment may incur penalties in another. Mesophilic bacteria have adapted to life in intermediate conditions where several dimensions of the cellular environment can readily cause stress, such as excessively high or low temperatures, acidity, nutrients, osmotic pressure, chemical concentrations, and so on. In the lifecycle of an enteric bacterium, drastic changes arise within and between the host and *in natura* conditions.

One such stress is antibiotics. The widespread use of antibiotics to treat livestock for enhanced meat production as well as the rise of antibiotic-resistant bacterial strains in medical contexts lends the question practical importance. Genetic resistance may also have an important relationship with transient antibiotic tolerance: persistence may allow resistance to evolve more quickly (3, 4), though this hypothesis has been contested. Multiple mechanisms induce persister formation (5–9), including starvation or loss of metabolic activity (10–14).

**Figure 1.**
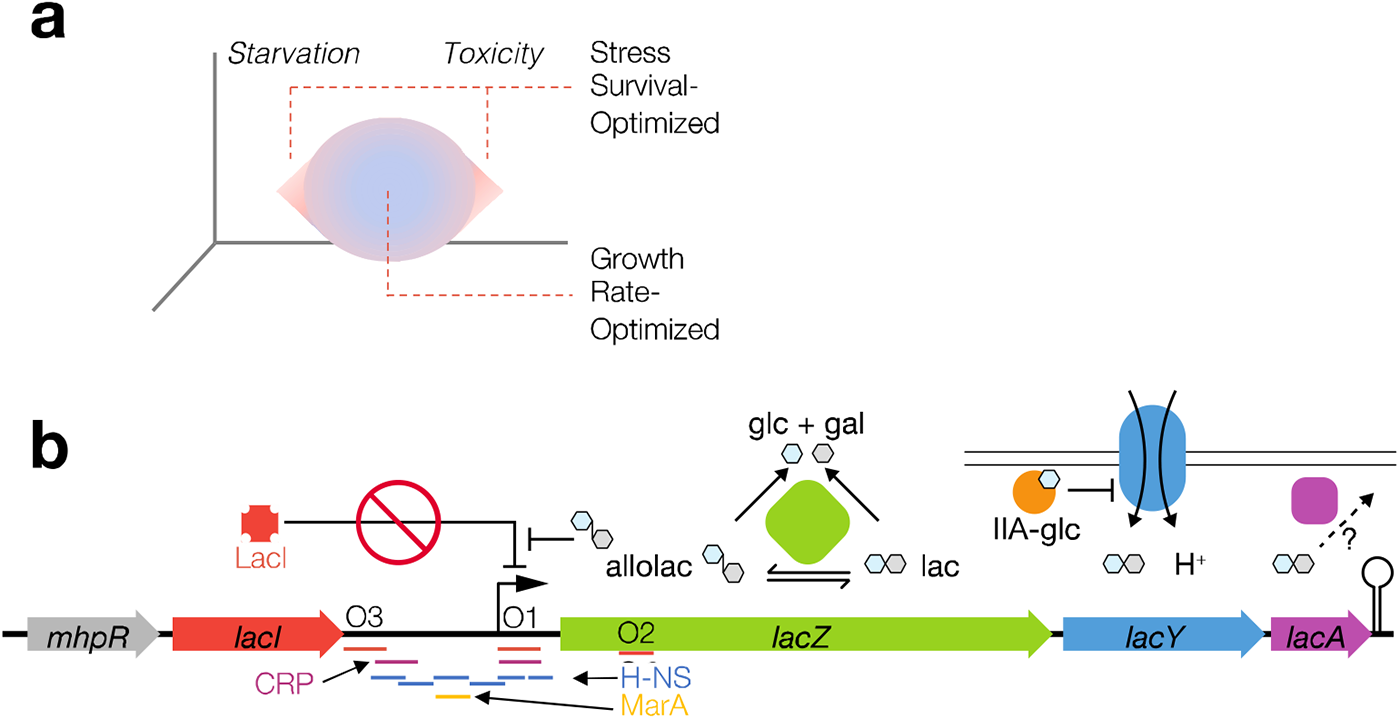
Fitness in a mesophilic microbe and the *lac* operon of *Escherichia coli* specifically. **a.** There is an envelope of survivable conditions in mesophilic bacteria. The volume labeled “Growth Rate-Optimized” denotes conditions to which the microbes are well-adapted for fast growth. In the peripheral volume, “Stress Survival-Optimized” (e.g., the stringent response and bet-hedging), the combination of environment and response improves colony survival but not growth rate. **b.** The mechanism to perturb fitness in this study is titration of lactose as a sole carbon source in minimal media. Lactose processing involves intrinsic physiological costs and may cause downstream toxicity in certain conditions via the Leloir pathway (galactose processing; not shown).

We previously demonstrated that excessively high or low lactose concentrations (as a sole carbon source) can predispose populations of bacteria derived from a strain of *Escherichia coli*, B REL606, to lowered death rates in antibiotics (15). Varying the concentration of lactose in a minimal medium drives a non-monotonic relationship with the exponential growth rate in culture, with a fast-growth plateau at intermediate concentrations (15). Lactose has established toxic effects on *E. coli* cultures, often attributed to membrane depolarization via excessive proton symport with lactose through the permease LacY, a member of the major facilitator superfamily of permeases (16–18). In B REL606, toxic lactose levels create a heterogeneous population dynamic with a chance of fast-growing cells to enter growth arrest, yet both starving and toxified cultures exhibit increased stress tolerance (15, 19; Fig. 2). Growth-arrested cells represent a persister-prone subpopulation in both conditions such that the toxified culture has the effect of a bet-hedging dynamic with average population growth rate sacrificed in favor of stress tolerance in a minority of the population. Does the global transcriptional program of stress responses have overall similarities between these conditions, or are key alternate pathways activated? The mechanisms for these conflicting stresses to attain a similar phenotype are unknown.

**Figure 2.**
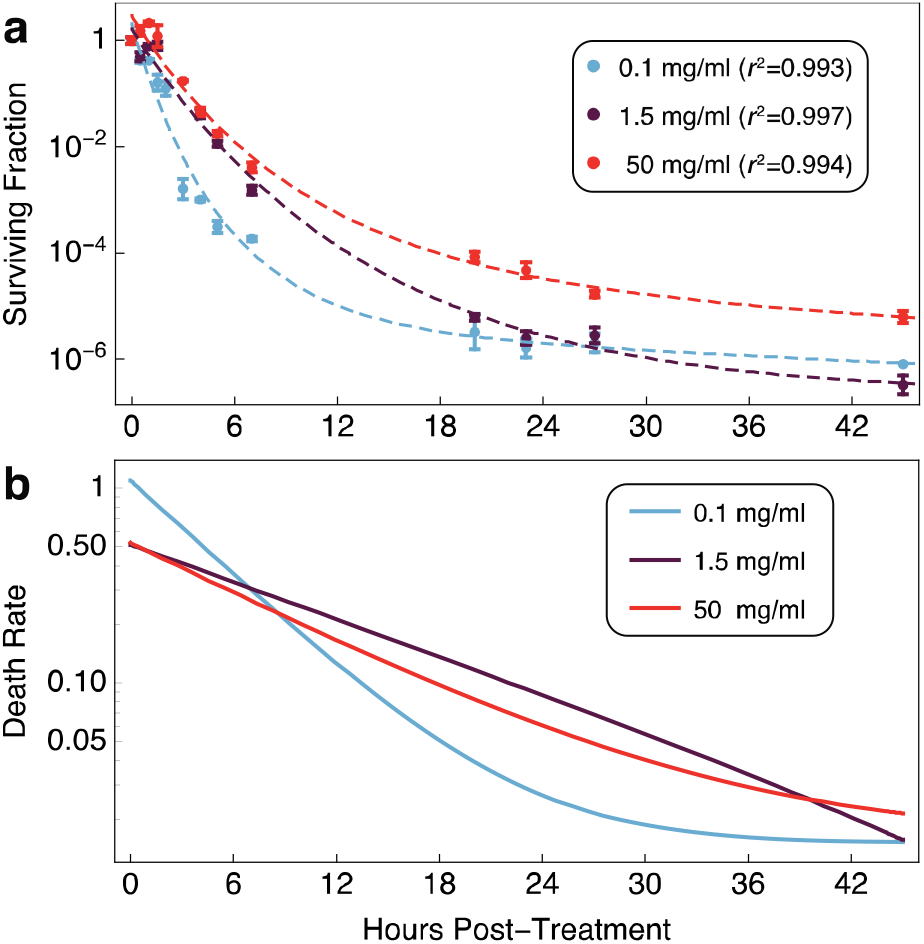
The killing rate of an *E. coli* B strain in ampicillin is lowest in starving and toxified culture conditions. **a.** Surviving fractions in low, intermediate, and high lactose conditions after ampicillin treatment (100 mg/ml) during mid-exponential phase (mean + standard deviation, *N* = 3; final point in 1.5 mg/ml, *N* = 1). Data from (2) were fit to a mixed linear-exponential model *y == a t + b e^-g t^ +c* with *r*^2^ as reported in the figure. **b.** Time derivative 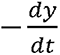 of the statistical model parameterized for each fit in panel **a** shows a lowered killing rate for both starving and toxified cultures between approximately 7 and 40 hours post-treatment. Model fit parameters: *(dose): (a, b, c, g)* (0.1): (-0.0149675, 5.71624, -5.40263, 0.189278) (1.5): (0.00427562, 7.13623, -6.91445, 0.072367) (50): (-0.0161676, 4.99618, -4.53027, 0.101607)

We interrogated this system with the bulk (population-level) RNA-seq under the hypothesis that a common stress response is conserved between starvation and toxicity. While making use of bioinformatic, biochemical, and physiological data about specific mechanisms, we avoided presuming that any particular mechanism was at work beyond the proven phenomena arising in these culture conditions. We cultured *E. coli* in low, intermediate, and high concentrations of lactose minimal medium, grew them to mid-exponential phase, and harvested biomass from each culture for bulk RNA-seq. The resulting transcriptional profiles were subjected to differential expression analysis with the intermediate lactose concentration as the reference condition.

## 2. Results

### 2.1. Slowed killing rate in starving and toxified E. coli cultures

Re-analysis of our previously measured timecourse of exponential-phase *E. coli* treated with ampicillin demonstrates a lowered rate of death in starving and toxified cultures (Fig. 2). To confirm this, we fitted the data to a mixed linear-exponential statistical model in logarithmic coordinates on the *y* axis: *y = a_t_ +b_e_^-gt^+c*. All three parameters were fit with low error (*r*^2^ 0.99). Taking the time derivative of the statistical model revealed the estimated rate of killing for each culture condition (Fig. 2b): highest in non-stressful conditions. Thus, both the starving and toxified cultures are prone to produce higher levels of antibiotic tolerance than less stressful intermediate conditions.

### 2.2. Differential gene expression analysis reveals growth-mediated shifts in lac operon expression, toxin-antitoxin responses, and global regulator responses

To analyze gene expression profiles, we purified total RNA from early-mid-exponential cell culture in different lactose concentrations. We mapped sequencing reads to the reference genome *E. coli* B REL606 NC_012967.1 (20) with kallisto (21) and subsequently analyzed the count data using DESeq2 in R (22) with subsequent processing in Python. Setting the moderate lactose concentration (2.5 mg/ml) condition as the reference, we defined differentially expressed genes (DEGs) in *starvation* (lactose conc. 0.1 mg/ml) and *toxified* (lactose conc. 50 mg/ml) conditions as genes with an adjusted *p*-value (false detection rate; FDR) of below 0.05 and a log_2_ fold change (LFC) greater than 1 (Figure 3).

**Figure 3.**
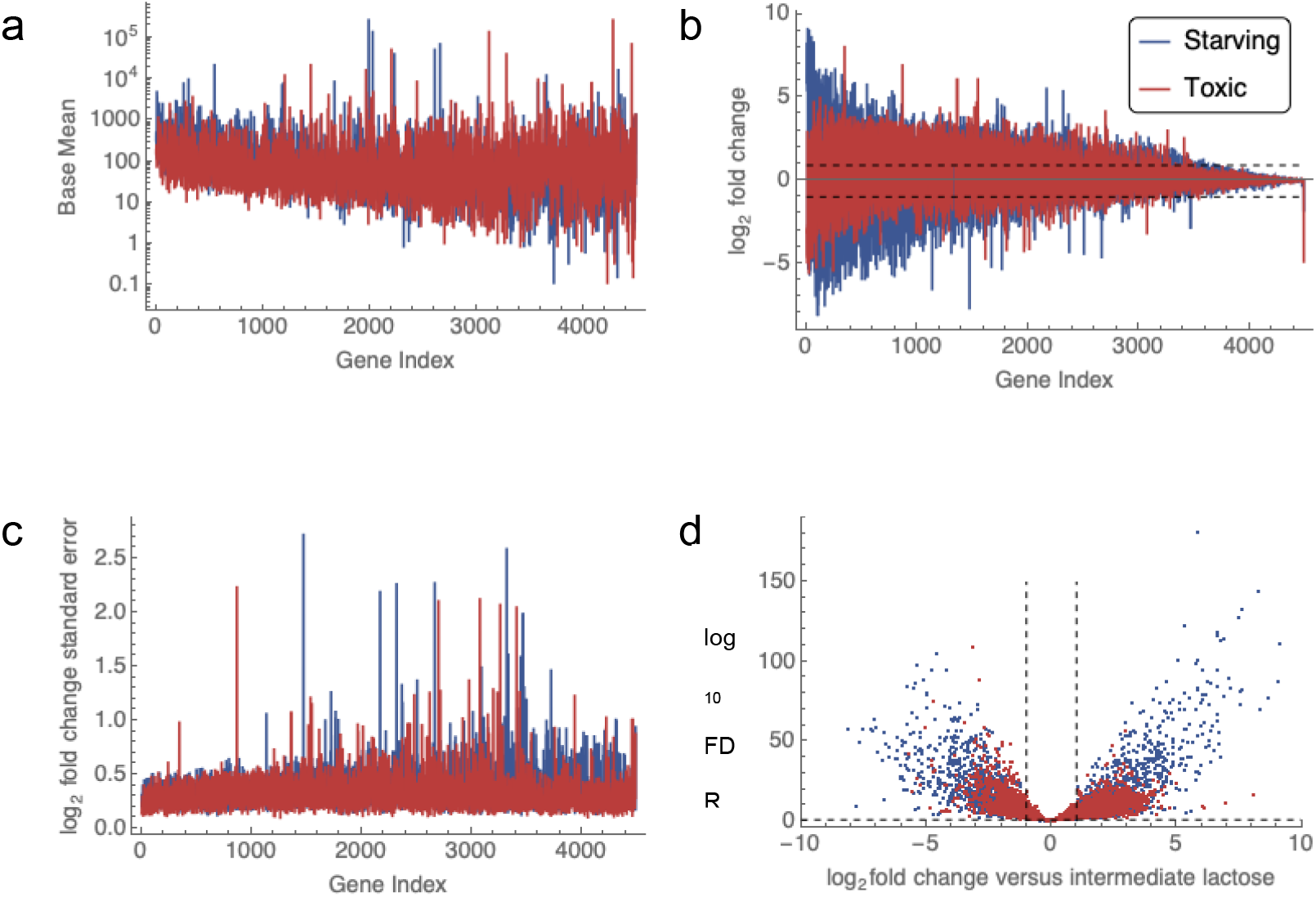
Comparison of differential expression profiles between starving and toxified cultures, both relative to a nominally lower-stress condition. **a.** Base mean number of reads. **b.** Mean fold change of each gene sorted by FDR from lowest to highest (Gene Index). Dashed lines indicate > 2-fold change in expression compared to intermediate-lactose cultures. **c.** Standard error of the fold change mean between biological replicates. N = 7 biological replicates for starving cultures, 9 for toxified cultures, and 9 for intermediate lactose cultures. **d.** Volcano plot reveals the extent of differential expression. Dashed lines indicate thresholds for meeting the criterion for significant differential expression: > 2-fold change and false detection rate (FDR) < 0.05.

In Figure 3a, the base mean is the normalized mean expression level for each gene in all replicates in the culture condition. The standard errors from the shrunken log_2_ fold change to corresponding maximum-likelihood estimates are well controlled (Figure 3c). They aligned well for a proper size factor calculation. Overlaying the differential expression log_2_ fold change shows that the starving cultures had wider log_2_ fold changes, suggesting a more extensive regulon, and perhaps more severe stress, compared to toxified cultures. This result is consistent with our previous results (15) and the model shown in Figure 1, where cells have a higher death rate under low lactose compared to toxic lactose conditions.

Out of 4490 genes in the genome, 1845 DEGs were upregulated and 1145 DEGs were downregulated in starvation, while 1755 DEGs were upregulated and 830 DEGs were downregulated in toxified cultures. Clustering of differentially expressed genes revealed that starving cells and toxified cultures have overlapping, but not identical, regulons (Figure 4a). Genes that were either upregulated or downregulated are color-coded points in Figure 4b. Many regulated genes trend toward equal differential expression (near the “line), but the data reiterate here the observation that starving cultures have a larger set of gene expression changes than toxified cultures^1^.

**Figure 4.**
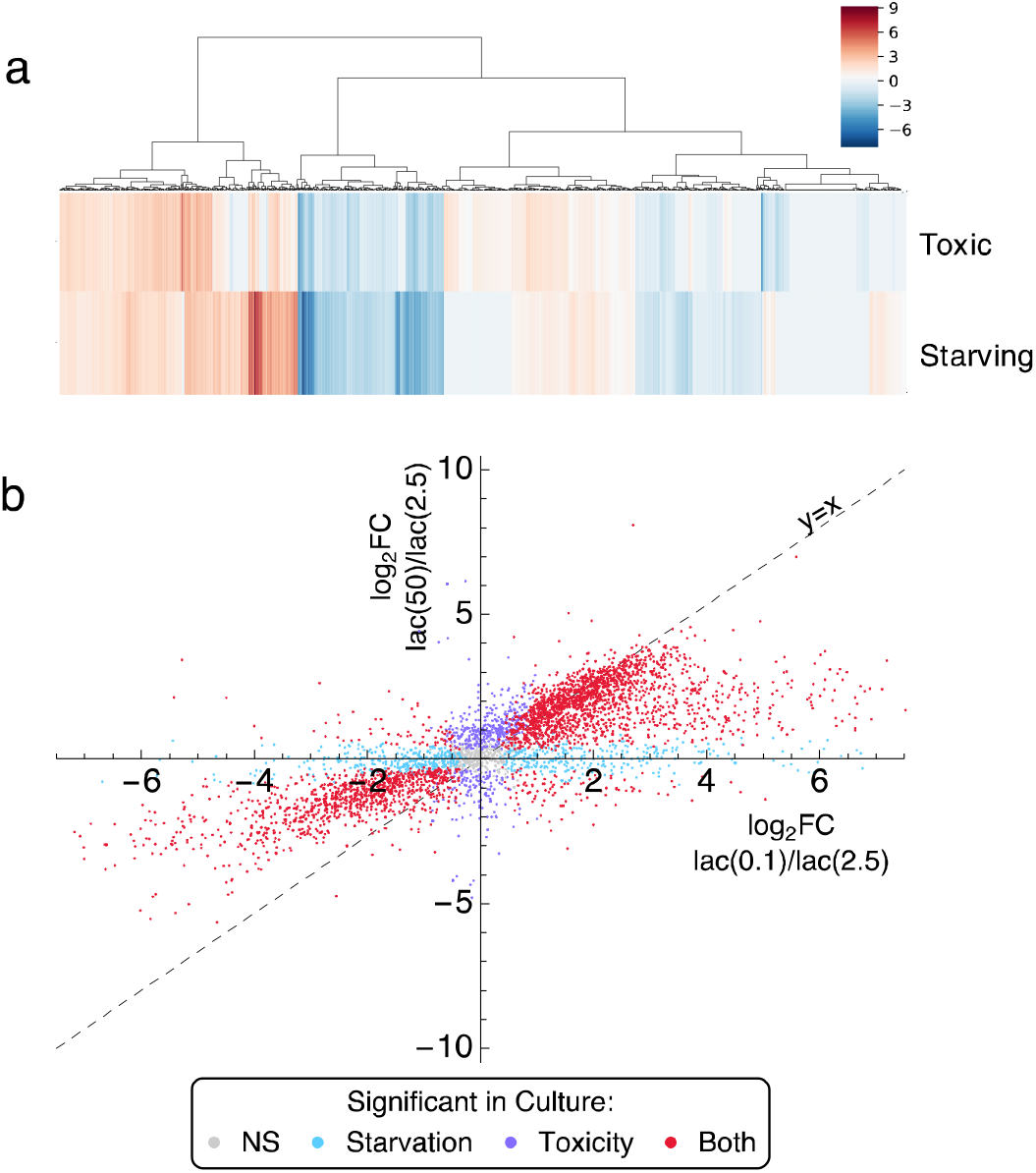
Fold change of genes in starving cultures versus toxified cultures relative to the intermediate baseline. **a.** DEGs clustering based on the log_2_ fold changes for each gene, where toxified cell profile behaves quite different from starved cells. **b.** Points are genes color-coded based on which conditions have significant fold change (FC). NS, non-significant.

Expression of the *lac* operon is a natural question in experiments where the quantity of lactose in cultures is varied. The *E. coli* strain used here lacks a functional lactose regulator (LacI), an intentional choice to disentangle the downstream consequences of lactose processing without the added complication of *lac* operon gene regulation. Because the *lacI* gene has a point mutation, detected transcripts encoding LacI were non-functional (Figure 5). Growth in lactose minimal medium requires *lac* operon expression in any case (Figure 1b). Any apparent differential expression of *lac* operon gene products is attributable to the following remaining factors (23): silencing by H-NS and MarA, activation by CRP-cAMP, σ^70^-activated promoters, and effects on mRNA concentrations by differential loss from either growth or a shift in mRNA degradation. The particular strain we used also expresses a recombinant GFP constitutively, which we have previously exploited as an optical proxy for growth rate because slower-growing cells accumulate GFP and become brighter (15, 19).

Following the steps of lactose processing is informative to see the difference between molecular gene regulation and the consequences of fitness, growth, and selection on gene expression in culture. LacY is the lactose permease, transferring extracellular lactose across the inner membrane into the cytoplasm (Fig. 1b). The starved cells contain substantially increased expression of LacY mRNA, suggesting that surviving cells in the culture have increased lactose uptake. In toxified cultures, *lacY* is not differentially expressed, suggesting relaxed selection for high lactose permease levels. The gene *lacZ* encodes β galactosidase, the catabolic enzyme that directly assimilates lactose into the metabolic network. *lacZ* is downregulated in toxified cultures, which may reflect the fact that most cells in the toxified culture randomly avoided toxicity by lowered lactose assimilation. Galactose degradation (the Leloir pathway) is directly downstream of lactose degradation by β-galactosidase, and UDP-galactose and galactose-phosphate intermediates in the Leloir pathway can cause stress and reduce cell growth (24, 25). Differential expression of *lacA* has unclear functional consequences and may simply reflect co-expression with *lacZ* and *lacY* from the operon.

We next explored differential expression of global regulators of growth rate: toxin-antitoxin systems (Fig. 5b) and transcriptional regulators (Fig. 5c). With gene annotation matching (see Methods), we found a total of 15 toxin-antitoxin (TA) modules existing in the B REL606 strain of *E. coli*. Among these TA modules, starving cultures exhibited activation of 6, including *ygiT-mqsR*, YhfG-Fic, Xre-HipA, *yeeUV*, *ghoTS* and *hokE-lexA*. Interestingly, *ygiT-mqsR*, *yeeUV*, and *ghoTS* are also upregulated in toxified cultures, while the other 3 TA systems show upregulated toxin genes. Knockouts of the global regulators in Fig. 5c result in at least a 10-fold decrease in persister formation (26). In starving cultures, 6 global regulators out of 9 are downregulated, including stress response regulator genes *dnaKJ*, DNA protection related regulator genes *hupAB*, a folate-dependent enzyme inhibitor gene *ygfA*, and persister cell formation regulator gene *yigB*. Though the recombination-related regulator genes *ihfAB* are upregulated, suggesting a possible higher mutation rate, *dksA* is also upregulated; the latter’s product can promote DNA recombination repair (27). The product of *dksA* is also an rRNA repressor that mediates stress responses and inhibits DnaK. Thus, we observe a pattern of gene regulation with counterbalancing effects that may be consistent with phenotypic decisions being made post-translationally. In toxified cultures, *ihfB* and the chemotaxis signaling genes *cheZ* and *cheY* were upregulated.

**Figure 5.**
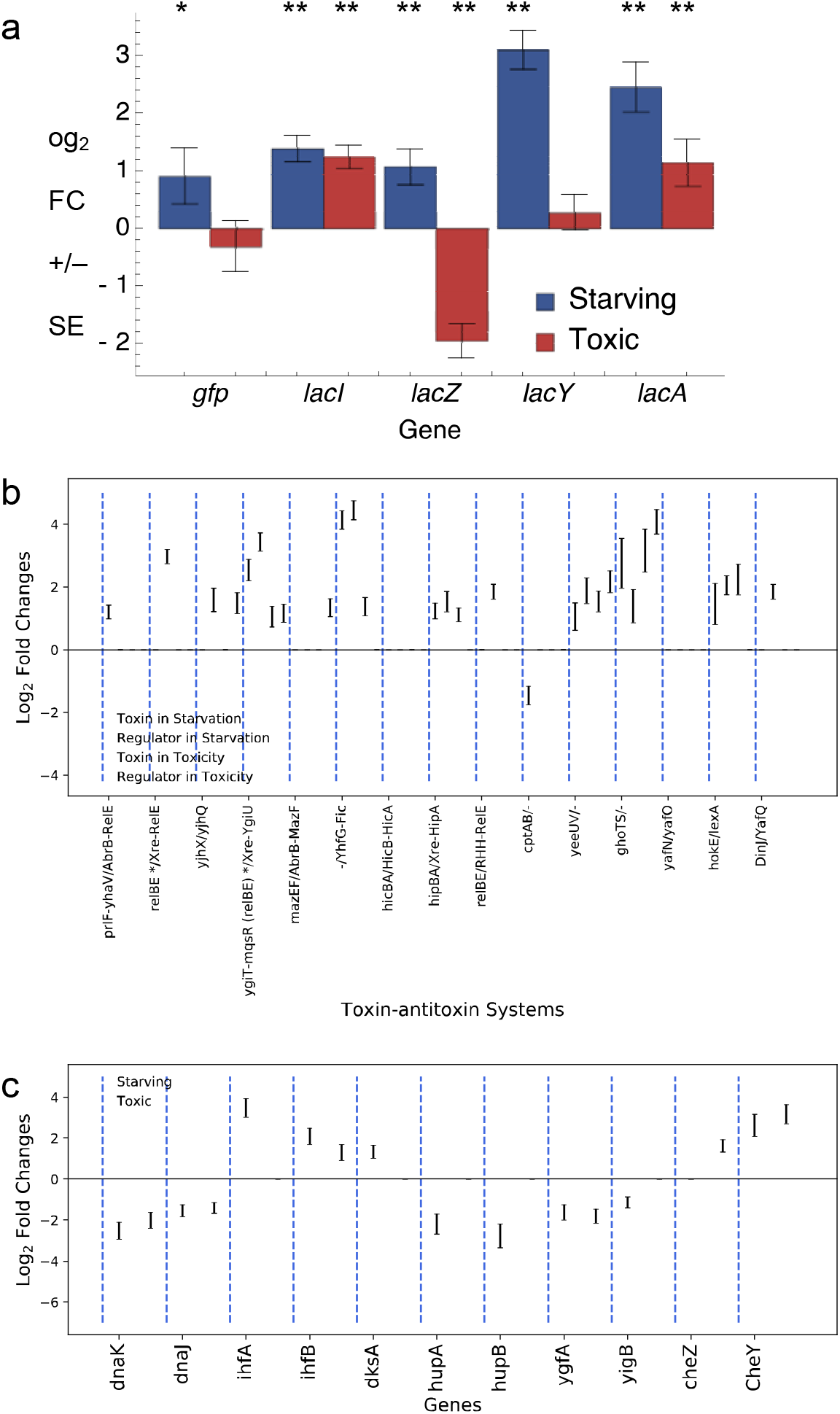
Differential expression log_2_ fold changes of reference genes in starving and toxified cultures establishes a baseline reference for persister-prone conditions. **a.** lactose operon activities. In this strain, the *lacI* gene is present but non-functional because of a mutation. As a result, the *lac* operon is expressed constitutively, as is the inserted *gfp* gene. FC, fold change. SEM, standard error of the mean. N = 7 for starving cultures, 8 for toxified cultures, and 8 for intermediate lactose cultures. **: FDR < 0.001; *: FDR < 0.05. **b.** Toxin-antitoxin systems. **c.** Global regulator genes and chemotaxis signal transduction related genes (*cheZ* and *cheY*).

### 2.3 Stress responses regulated by sigma factors

Transcription initiation for *E. coli* promoters requires sigma (σ) factors, subunits of RNA polymerase (RNAP) (28), making them global regulators of responsiveness to the environment. Individual promoters typically respond to specific σ factors. Six σ factors are known in *E. coli* B REL606. σ^28^ (RpoF/FliA), the flagellar synthesis sigma factor in *E. coli* strain K-12 MG1655, is missing in REL606 and σ^19^ (FecI), the ferric citrate sigma factor regulating iron transport and metabolism, is present but does not have any annotated gene regulatory roles in RegulonDB (29). To understand how σ factors regulate cellular responses to environmental stimuli, we explored their differential expression between lactose concentrations and that of their regulated genes as log_2_ fold changes (LFC2).

**Figure 6.**
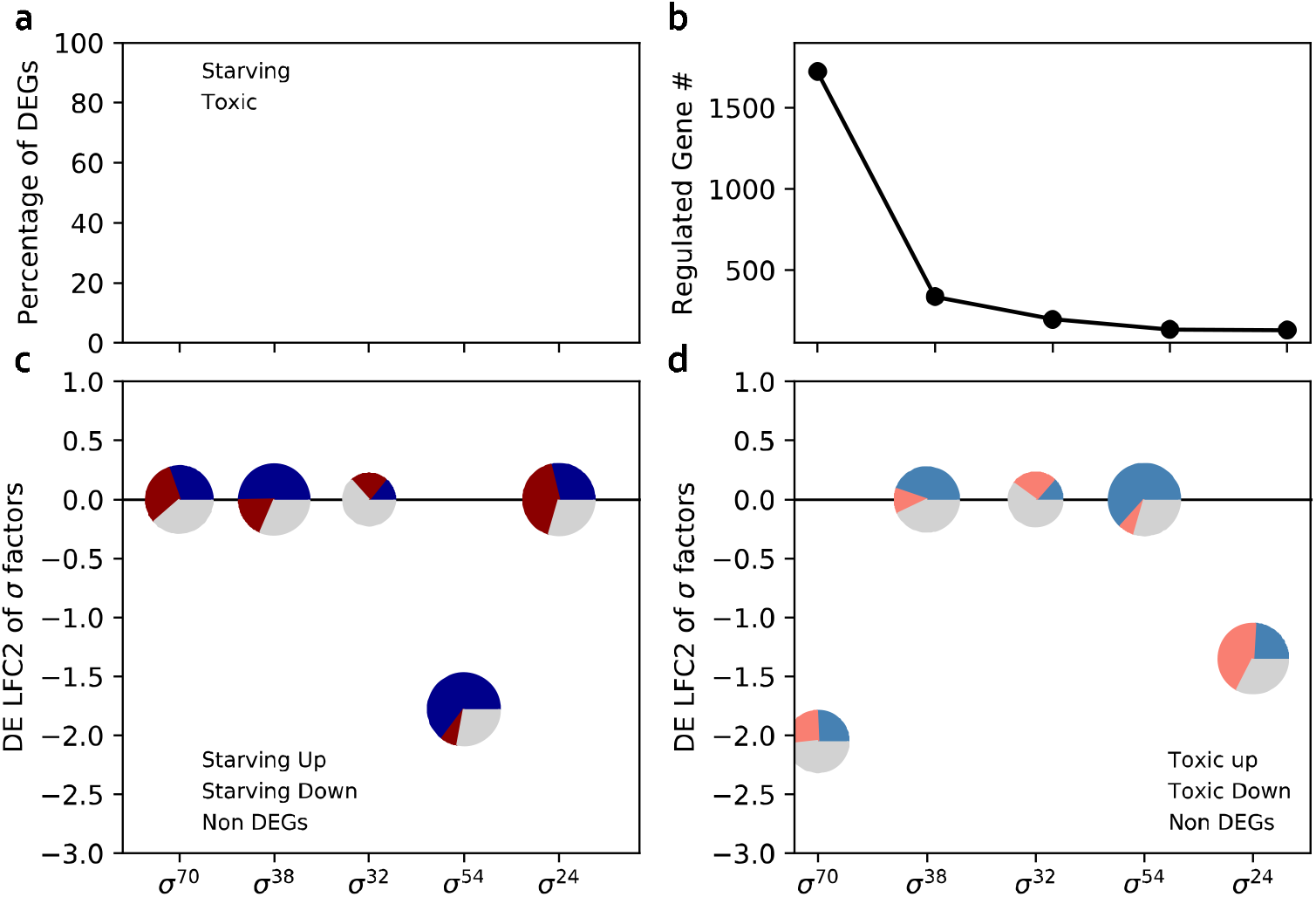
Regulation of σ factors differ in response to starvation and toxification. **a.** Percentage of differentially expressed genes (DEGs) in each σ factor regulon. **b.** The number (#) of genes annotated to be regulated by each σ factor. **c**, **d**. Differential expression (log_2_ fold change) of the σ factors and their regulons in **c.** starving and **d**. toxified cultures. The size of the pie chart reflects the DEG percentage regulated by each sigma factor. LFC2, log_2_ fold change.

Our results show a significant decrease in σ^70^(*rpoD*) in toxified, but not starving, cultures (Fig. 6c,d). This σ factor initiates transcription in a set of approximately 1723 genes involving cell proliferation-related behavior such as substrate uptake, DNA replication, membrane synthesis, and ribosome production (23). About half of the genes initiated by σ^70^ were downregulated in both toxified and starving conditions, suggesting an overlapping regulon between the stress responses in starved and toxified cultures.

σ^38^ (rpoS), the sigma factor associated with the stringent response, does not appear to be differentially expressed on average in starving or toxified cultures, yet a plurality of genes regulated by σ^38^ is upregulated in both conditions as compared to growth-optimized lactose concentrations. This observation is consistent with σ^38^ regulation arising from sigma factor competition, especially in toxified cultures. The ratio of σ^70^ to σ^38^ in toxified cultures is consistent with this model as well. (p)ppGpp is involved in both production and activity of σ^38^ (30). The (p)ppGpp synthases, *relA* and *spoT*, are downregulated in both starving and toxified cultures. This is not necessarily inconsistent with the presence of the stringent response, as (p)ppGpp is known to exhibit a transient increase early in the stringent response, possibly before we sampled the cultures for RNA-seq (31).

σ^54^ (rpoN) stimulates transcription initiation in 136 genes involving nutrient limitation such as nitrogen assimilation, substrate-specific transport systems, and utilization of alternative carbon and energy sources. Most genes in the σ^54^ regulon are downregulated in both starving and toxified cultures, despite σ^54^ itself being downregulated or not significantly differentially expressed in starving and toxified cultures. The best-characterized mechanism for σ^54^ interactions with σ^38^ is through the glutamate-dependent acid resistance (GDAR) system (*gadE, gadA, gadBC*) (32). σ^54^ is described as a repressor for GDAR system, while σ^38^ is annotated as an activator.

σ^24^, driving responses to heat shock and other stresses on the membrane and periplasmic proteins, is downregulated on average in toxified cultures but is not significantly regulated in starvation here. σ^24^ is required for transcription of *degP*, a gene coding for a protease that degrades abnormal proteins in the periplasm (33). Downregulation of σ^24^ may result in less DegP protease, causing accumulation of DegP targets. Such accumulation is associated with increased membrane resilience, which is relevant because one of the factors thought to stabilize this strain of *E. coli* in toxified cultures is resistance to osmotic stress caused by excessive lactose import (see Discussion).

In all, downregulation of σ^70^ may drive a stimulon associated with σ^38^- and σ^54^-compatible promoters. The majority of genes regulated by σ^38^ and σ^54^ encode stress responses in both nutrient scarcity and conditions of over-abundance. However, sigma factor log_2_ fold changes are not always consistent with the dominant trend of gene differential expression, which we interpret to mean that post-translational interactions drive part of the response and that additional regulatory factors play a role in the responses of *E. coli* to starving and toxified conditions.

As starvation and toxicity are both intertwined with metabolism, we performed metabolic pathway gene set enrichment analysis in peripheral (Section 2.4) and central (Section 2.5) metabolism. We used the normalized enrichment score (NES), a statistic that reflects the degree that a pathway is overrepresented at the top or bottom of a ranked list of genes (Methods Section 4.5). An overview of our pathway enrichment analysis is presented in Figure S2.

### 2.4 Peripheral metabolic pathway gene set enrichment analysis

#### 2.4.1 Pathway regulatory similarities between starving and toxified cultures

Gene set enrichment analysis calculates trends for defined gene sets. The similarity of pathway regulation was calculated using rank correlations (Spearman’s ρ) with differential gene expression profiles (see Methods Section 4.5.1).

Figure S3 shows the correlation between pathway regulation in starvation and toxicity. Each node in Figure S3 represents a pathway, with a total of 362 pathways annotated in *E. coli* B REL606. Nodes are connected by metabolite edges as annotated in EcoCyc. We found that most metabolic pathways are either similarly regulated or have no correlation between treatments. We observed conserved regulatory pathways for persister-prone phenotypes, including central metabolic pathways such as the pentose phosphate pathway, chorismate biosynthesis I, glycolysis I, and the superpathway of glyoxylate bypass and TCA. Glycolysis is the hub node for the pathway network with similarly regulated pathways linking to glycolysis with degree 1 node distances. Statistically significant oppositely regulated pathways are rare. The only oppositely regulated pathway with statistical significance is tetrahydrofolate (FH_4_) biosynthesis, which is upregulated in starvation but downregulated in toxicity. FH_4_ biosynthesis produces vitamin B9 (folic acid), a cofactor leading to the biosynthesis of methionine, purines, thymidylate and pantothenate.

#### 2.4.2. Common enriched pathways in persister-prone phenotypes

Among 362 enriched pathways, 44 were upregulated in starvation and toxification, 69 were downregulated in starvation and toxification, 14 were uniquely upregulated in starvation, 31 pathways were uniquely downregulated in starvation, 12 pathways were uniquely upregulated in toxification, and 25 pathways were uniquely downregulated in toxification. Detailed pathway enrichment analysis results can be found in Supplementary Tables S1-S6. Figure 7 shows the pathway hierarchy structure based on EcoCyc annotations. The directed edges point to the parent pathway which is often consisted of multiple pathways to form superpathways. In Figure 7 we mapped the normalized enrichment score calculated with FGSEA (34) onto the pathway hierarchy graph. The result is many clustered enriched pathways. The top 3 clusters in Figure 7 are (1) the superpathway of chorismate metabolism, (2) amino acid biosynthesis, and (3) the superpathway of histidine, purine, and pyrimidine biosynthesis.

**Figure 7a.**
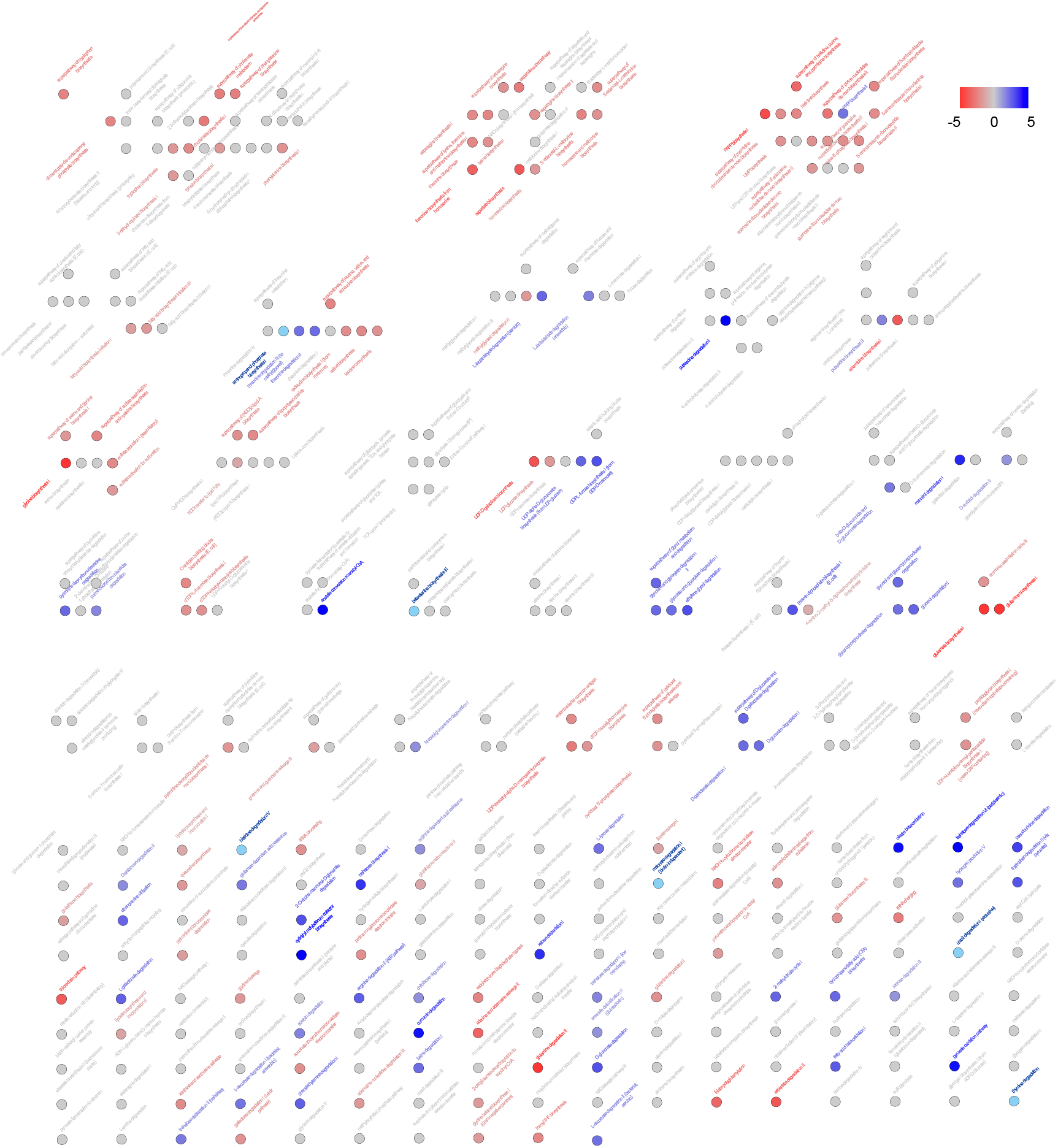
Pathway enrichment score graph representation for pathway ontology in starvation condition. The upregulated pathways are shown in blue, and downregulated pathways are shown in red. The pathway ontology annotation is from EcoCyc The pathway ontology annotation is from EcoCyc (1).

**Figure 7b.**
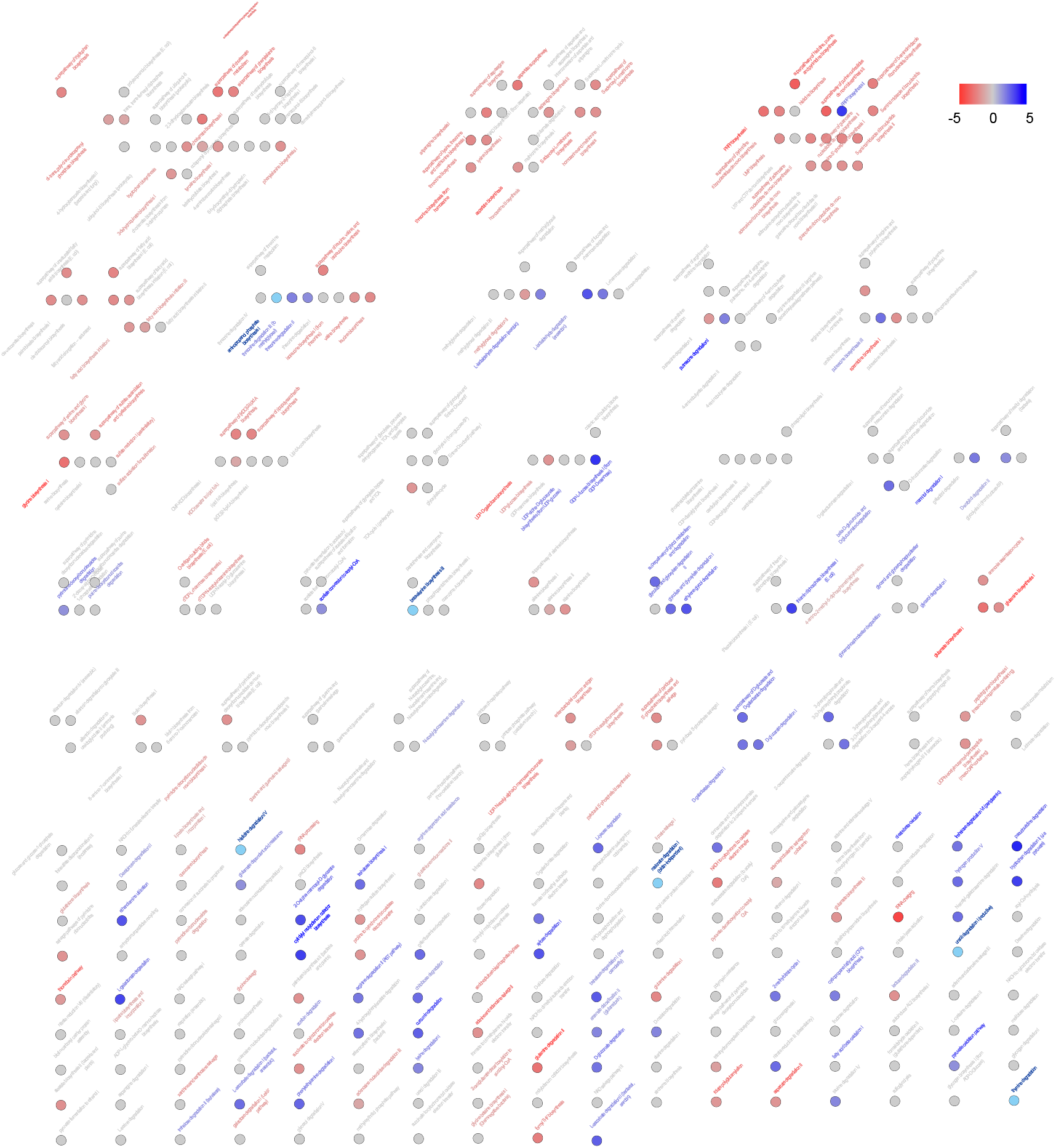
Pathway enrichment analysis graph representation for pathway ontology in toxic condition. The upregulated pathways are shown in blue, and downregulated pathways are shown in red. The pathway ontology annotation is from EcoCyc (1).

Chorismate is the principal precursor for the aromatic amino acids, such as tryptophan, tyrosine, and phenylalanine (35). Downregulation of chorismate biosynthesis can introduce aromatic amino acid starvation (this may occur in both toxication and starvation according to the enrichment profile). Menaquinol-8, ubiquinol-8, tetrahydrofolate biosynthesis, and enterobactin biosynthesis are not differentially regulated compared to the cultures with intermediate lactose concentration (chorismate metabolism provides essential compounds for those pathways). The quinone pool is essential for *E. coli* adapting to different oxidative conditions and maintaining proper redox- and phosphoryl-transfer reactions to form the core of cellular energetics (36). Biosynthesis of ubiquinone is reported to accumulate pathway intermediates (37, 38) with the effect of improving osmotic stress-tolerance.

We found that 17 out of 20 amino acid biosynthesis pathways are downregulated in persister-prone phenotypes accompanying upregulated amino acid degradation pathways (Table S2.). Alanine and arginine biosynthesis are uniquely downregulated in toxification, while asparagine is uniquely downregulated in starvation, implying lower NAD synthesis (39) and likely resulting in lower energy levels. Amino acid starvation can lead to accumulation of uncharged tRNAs that enter the ribosomal A site, halting translation (40). We observe downregulated tRNA charging in both toxicity and starvation. The pyrimidine, purine and pyridine nucleotide synthesis-related pathways are downregulated in both starvation and toxicity except for phosphoribosylpyrophosphate (PRPP) biosynthesis II, which provides PRPP as a pivotal metabolic precursor to pyrimidine, purine, and pyridine nucleotide synthesis. Other downregulated pathways include sulfate assimilation downstream of cysteine biosynthesis, lipid A, lipopolysaccharide (LPS) and peptidoglycan biosynthesis, spermidine biosynthesis, UDP-glucose biosynthesis, and cytochrome b_o_ oxidase electron transfer pathways from proline succinate.

Upregulated pathways have a sparser network structure (Table S1). Among 44 commonly upregulated pathways, 30 are degradation pathways targeting amino acids, other carbon sources, and electron transfer related metabolites, such as L-ascorbate (vitamin C) and putrescine. Broadly upregulated nutrient assimilation pathways reflect supervening carbon source uptake and balancing energetic electron transfer. Other upregulated pathways enhance cellular fitness through energy metabolism, overflow metabolism, and membrane component regulation. In both persister-prone conditions of this study, the pyruvate oxidation pathway is upregulated; this pathway generates the transmembrane potential for constructing respiratory chains consisting of pyruvate oxidase, ubiquinone-8, and the cytochrome *bd* complex (41). Thiamine diphosphate biosynthesis I produces cofactor vitamin B1, which plays a fundamental role in energy metabolism. Trehalose pathway upregulation affects the osmotic stress response, notably present in both stress conditions. In response to acetate and osmotic pressure, the arsenate efflux pump, GDAR system, and aerobic utilization of acetate are upregulated. GDP-L-fucose biosynthesis I produces LPS components in the membrane, and membrane structure may be further stabilized by enrichment in putrescine biosynthesis III, which is the precursor for spermidine biosynthesis.

#### 2.4.3. Uniquely enriched pathways in starvation

Pathways uniquely upregulated in starvation (Table S3) include nutrient assimilation from metabolites mannitol, D-arabinose, acetoin, glycerol, glycolate, glyoxylate, trehalose, and fatty acids. The arginine-dependent acid resistance pathway was also observed.

Though downregulation (Table S4) of the Leloir pathway may be beneficial to reduce UDP-glucose induced toxification, it may also limit production of α-D-glucopyranose-1-phosphate, a precursor for synthesis of the outer membrane LPS component O-antigen. Despite O-antigen synthesis precursor UDP-α D-glucuronate biosynthesis being upregulated, biosynthesis of O-antigen building blocks from precursors including α-D-glucopyranose-1-phosphate and sugars L-rhamnose, and dTDP-*N*-acetylyiosamine are specifically downregulated in starvation.

Downregulation of glutathione biosynthesis and redox reaction III may play a role in redox balance and defense against reactive oxygen species.

Several other pathways uniquely downregulated in starvation suggest a trend toward energy conservation: downregulation of lipoate metabolism, the pentose phosphate bypass, sedoheptulose bisphosphate metabolism, thiamine precursor biosynthesis, nucleotide precursor biosynthesis, and several acyl-CoA derivatives.

Finally, asparagine biosynthesis pathways are uniquely downregulated in starvation, with unclear functional significance.

#### 2.4.4. Pathways specifically enriched in toxicity (Table S5)

Two pathways forming the *E. coli* anaerobic respiratory chain are upregulated: the formate-to dimethyl sulfoxide electron transfer pathway and nitrate reduction III, implying increased proton-motive force across the cytoplasmic membrane (42–45). We observed upregulation of taurine degradation IV, which provides alternative sulfur under sulfate starvation induced by cysteine starvation (we note that our culture conditions are similar to those often used with this *E. coli* strain in the LTEE: Davis minimal medium containing supplemented lactose and thiamine (46) without amino acid supplementation). Several sugar degradation pathways are uniquely upregulated in toxicity, suggesting a possible carbon source alternative for *N*-acetyl-galactosamine (GalNAc), D-galactosamine (GalN), D-malate, L-rhamnose, and galactitol. Other enriched pathways are mostly fermentation metabolic pathways.

The superpathway of fatty acid biosynthesis initiation is uniquely downregulated in toxified conditions (Table S6). Related pathways such as fatty acid elongation, palmitoleate biosynthesis, and *cis*-vaccenate biosynthesis are also uniquely downregulated. Fatty acids are key building blocks for phospholipid components of cell membranes and are determinants of intracellular communication where palmitoleate is a common unsaturated fatty acid. Farnesyl pyrophosphate (FPP) biosynthesis is downregulated, possibly implying less membrane attachment in posttranslational modifications.

### 2.5. Gene set enrichment analysis of central metabolism

To complete the analysis of regulated metabolic pathways in starvation and toxicity, we mapped differential expression to central metabolism (Fig. 9). As the culture conditions in this study demand that the initial entry to central metabolism occurs via lactose degradation, we define central metabolism herein to be composed of the lactose degradation pathway, pentose phosphate pathway, glycolysis, Entner Doudoroff shunt, TCA cycle, and the glyoxylate bypass. As galactose is a direct product of lactose degradation, we included the Leloir pathway (galactose degradation I) as well.

In starvation, the glycolysis pathway is mostly downregulated except for one gene encoding a phosphate transfer reaction. Components of glycolysis are more consistently upregulated in toxicity. The Entner-Doudoroff shunt linking glycolysis to the TCA cycle is regulated oppositely between starving and toxified cultures. The shunt is upregulated in starvation, possibly providing paths to alternative metabolites. Though the TCA cycle is largely downregulated in both conditions, the glyoxylate cycle is upregulated by various degrees in both starvation and toxicity. The glyoxylate cycle is a bypass for the TCA cycle to skip steps that remove CO_2_, thus conserving carbon intermediates during growth (47–49). The pathway is repressed during growth on glucose, and induced by growth on acetate, the byproduct of overflow metabolism.

**Figure 9.**
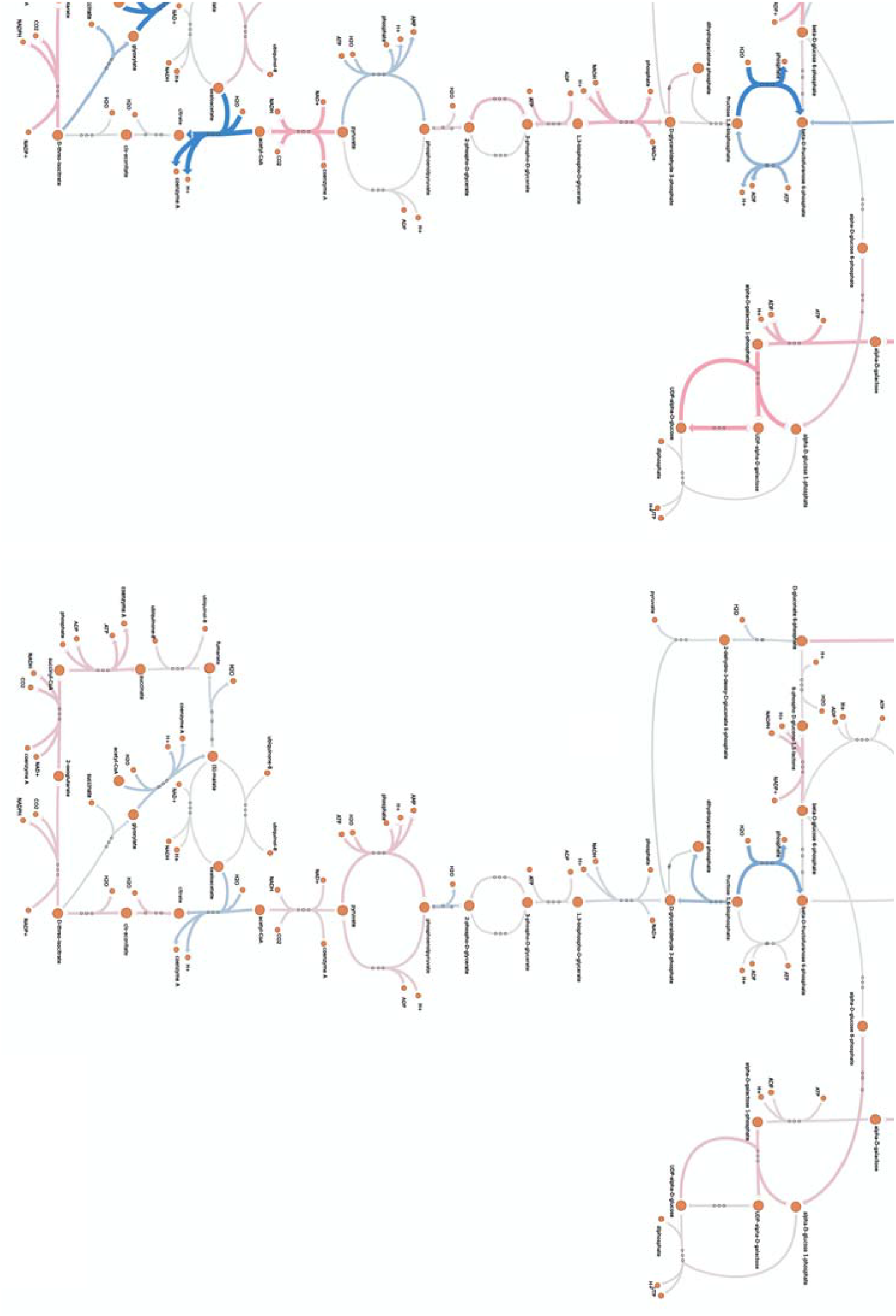
Differential expression (log_2_ fold changes) of genes encoding central metabolism in **a** starving and **b** toxified cultures.

## 3. Discussion

Variation in fitness-relevant environmental properties affects cellular gene expression patterns in quantitative and qualitative ways. Our previous demonstration that an excess of a required nutrient drives the formation of antibiotic tolerance (15) provided an opportunity to re-evaluate the nature of integrated microbial stress responses. This phenomenon appears to arise from the robust nature of the cell wall in B strains of *E. coli*, which permits survival in adverse osmotic conditions (50). We were specifically able to enrich antibiotic-tolerant cells in conditions both below and in excess of optimal concentrations for lactose as the sole carbon source (Fig. 1). We created such conditions in batch culture to test hypotheses about the nature of antibiotic tolerance in *E. coli*: what kind of signal may predispose cells to stress tolerance, and do conditions that could be described as “opposite” of each other from osmotic, nutrient concentration, and media toxicity perspectives induce similar or largely different responses?

We conceptualized analysis of the stress response by analogy to a statistical model, with linearly independent (normalized) stress response components ξ*_i_* and interactions ξ*_ij_* between components ξ*_i_* and ξ*_j_* such that a given response *i* is independent from *j* if ξ*_ij_* = 0. The integrated response can then be characterized by 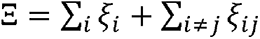 (eliding coefficients because we assume normalization between the components). The integrated response Ξ is a high-dimensional vector describing the phenotype. Independence between components may arise in some contexts and not others – we did not presume to comprehensively describe this space in our study, but rather to sample from it for exploration of the specific nature of stress responses. Nonzero interaction terms may arise from several sources, including pleiotropic effects (e.g., temperature or growth rate), co-expression (operon membership, transcriptional co-regulation, etc), and catalysis or synthesis of metabolites with broad effects (amino acids, nucleotides, tRNAs, etc).

Single cell dynamics play an important role in physiology and have a strong effect on lactose-toxified cultures. In the *E. coli* strain used in this study, a threshold drives toxicity to arrest growth in a subset of cells while subthreshold cells maintain growth. Thus, the results presented here represent an average across the subthreshold growing cells and the growth-arrested toxified subpopulation. The subthreshold subpopulation comprises approximately 10 percent of the entire population (15). This quantity arises from exponential growth of sub-threshold cells causing them to quickly overtake the population.

We exploited extensive mechanistic knowledge about the direct effects of pathways to interpret transcriptional signatures. Targeted experiments to further test other aspects of responses, such as post-translational effects, are beyond the scope of this study. Our results recapitulated previous interpretations of pathway analysis in stress but added crucial new results that broaden some, and narrow other, key interpretations. This study focused on the nature of the average gene expression in cultures known to be enriched for persister cells, but with no specific selection for them. In a following study, we will present an analysis of persister cells in the starving and toxified conditions that survived antibiotic treatment.

### 3.1. Lactose toxicity-induced persistence may arise via a combination of overflow metabolism and Leloir pathway intermediates

One hypothesis for non-starving persistence is that metabolic toxicity is induced by critical proteomic concentrations. Galactose degradation I (the Leloir pathway) is directly downstream of lactose degradation by β-galactosidase and may accumulate UDP-galactose and galactose-phosphate intermediates causing stress and loss of growth. With high metabolic rates, *E. coli* (and virtually every other species) undergoes a metabolic shift from primarily aerobic metabolism to incomplete oxidation of metabolites, including ATP synthesis (51). The cause seems to be linked to proteomic optimization, as anaerobic ATP synthesis requires a smaller fraction of the proteome to synthesize an equivalent amount of ATP at the cost of more sugar (52). The smaller proteome allows for higher growth rates due to the reduced size of the necessary metabolome, allowing more transcriptional/translational machinery to be devoted to ribosome synthesis (52, 53).

We found that toxic culture conditions that produce persister cells have apparent utilization of the Entner-Doudoroff shunt, which connects pyruvate to phosphoenolpyruvate, thus linking to the glyoxylate cycle and downregulation of the enzymes responsible for oxaloacetate and acetyl-CoA entering the citric acid cycle, consistent with cells undergoing overflow metabolism. As the only carbon source initially present in the medium is lactose, GalE fluctuation may lead to UDP-Galactose toxicity (24), and indeed, *galE* is downregulated in high-lactose antibiotic-tolerant cells, but not significantly differentially expressed in untreated cultures.

### 3.2. Different phenotypes may arise from sigma factor competition

Regulation of stress responses in bacteria is generally considered to be robust to a variety of conditions. Despite being exposed to opposite stresses, we showed that differential expression of genes in starvation and toxicity is positively correlated. This may be due to leaky expression of genes in the different loci on the genome or generalized stress responses. As expected, regulated differentially expressed genes have wider fold change distributions compared to constitutively expressed genes and more genes are regulated in starvation than toxicity. We found that the global proliferation regulator σ^70^ is downregulated in toxification, potentially reducing *lac* operon expression independently of LacI. As genes regulated by σ^38^ are mostly upregulated in starving and toxified cultures, our results are consistent with the sigma factor balance leaning toward σ^38^. The nutrient limitation-responsive sigma factor σ^54^ is downregulated in starving cells, aiding glutamate-dependent acid resistance (GDAR). In toxified conditions, downregulation of σ^70^ is drastic, moving the sigma factor competition balance towards σ^38^. Though σ^54^ is not downregulated, GDAR is again upregulated in toxified cells. Thus, nutrient-poor and nutrient-rich conditions both stress the cells with clear regulatory responses that overlap.

### 3.3 Non-transcriptional and undetected factors at play in stress-responsive gene expression

Survivorship bias, as opposed to targeted transcriptional regulation, may play a role in the observed transcriptome profiles obtained from stressed cells. Noisy gene expression dynamics creates a distribution of cell states, including some in sub-optimal transcriptional states. With exponential population growth, fitter cells quickly overtake the population and dominate the measured transcriptional signal. Furthermore, it was impractical to examine every possible transcriptional function. Some that we did not emphasize, such as prophage induction, may play an uncharcterized role in distinguishing responses to divergent stresses. Thus, studies of subpopulations and deeper targeted analysis may further clarify the he overall similarities between divergent stresses.

**Figure 10.**
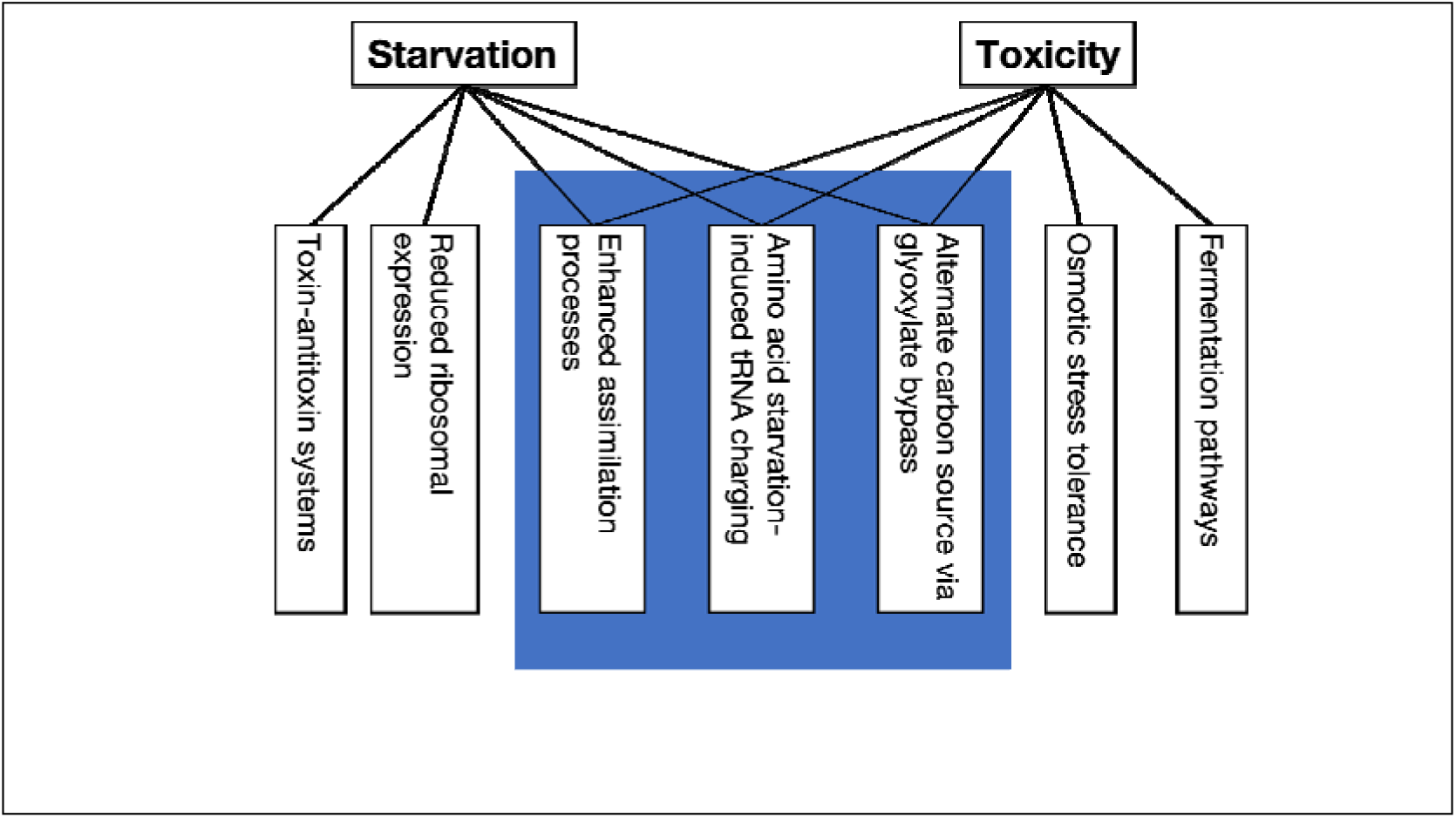
Overlapping and condition-specific stimulons in divergent stress responses of an *E. coli* strain.

### 3.4 Summary

The relationship between bacterial responses to divergent stresses contains four primary components: (*i*-*ii*) specific responses to each of the two stresses, (*iii*) a common stress-responsive regulon, and nd (*iv*) noise or functionally irrelevant responses. Overlapping responses include transcripts relating to nutrient assimilation, tRNA charging, and utilization of the glyoxylate bypass while condition-specific responses make sense in terms of unique properties of the stress in question (Fig. 10).

## 4. Materials and Methods

### 4.1. Persister Enriched RNA-Seq Experiments

*E. coli* B REL606 *lacI*^-^ transformed with Tn7::PlacO1GFP(KanR) was inoculated in LB medium from a -80 C bacterial stock and grown for 16 hours in a 37 C shaking incubator. The LB culture was then resuspended (1:1000) into 5mL of Davis Minimal medium (DM; Difco) supplemented with thiamine and one of three lactose concentrations (0.1 mg/mL, 2.5 mg/mL, and 50 mg/mL). The cultures were allowed to acclimatize for 24 hours before being resuspended (1:1000) into 5mL of the same Davis Minimal medium and lactose concentration. Cultures were grown long enough to provide enough biomass for RNA-seq after antibiotic treatment (8 hours in 2.5 mg/mL lactose, 10 hours in 50 mg/mL lactose, and 12 hours in 0.1 mg/mL lactose). After the initial growth phase, 1.5 mL cell culture aliquots were subjected to RNA isolation according to the following procedure.

Cell cultures were pelleted in a microcentrifuge (10,000 G for 2 minutes), washed in PBS buffer twice, resuspended in 500 μL of RNA-Later (ThermoFisher) and stored in at -20 C for up to one week. RNA isolation was performed using Direct-Zol (Zymo) and TRIzol reagent (ThermoFisher) and stored in a -80 C freezer overnight. Isolated RNA was ribo-depleted using RiboZero (Illumina) with ethanol washing to precipitate the RNA. Library preparation was completed using NEBNext Ultra II Directional RNA Library Prep Kit for Illumina (New England Biolabs) and sequenced using MiSeq v3 Paired End 150 bp (Illumina). Data are available from NCBI project number PRJNA938933.

### 4.2. RNA-seq sequence alignment and genome annotation with Ecocyc and RegulonDB

RNA transcript quantification was performed using kallisto (21) with reference genome NC_012967.1 (20) and 10 bootstrap samples. Functional interpretations used Ecocyc (Ver. 23.1) (54) and RegulonDB (55). Lacking extensive annotation of gene regulation in REL606, we used the K-12 MG1655 strain annotation based on gene names and gene product similarities.

### 4.3. Differential Expression Analysis

We used R package DESeq2 (56) for gene differential expression analysis. The RNA transcription quantification data were first clustered to isolate outlier replicates using principal component analysis (PCA) (Figure S2). Two samples with a high number of missing transcripts were dropped from subsequent analysis. To confirm reproducible outcomes of sample treatment, hierarchical clustering using the Wald significance test was performed on all samples; sample treatment was retrieved perfectly (Figure S1).

The DESeq2 pipeline includes size factor estimation, dispersion estimation, and DEG tests. Low count RNA quantifications are noisy and may decrease the sensitivity of DEGs detection (57). Size factors were calculated with a subset of control genes: non-regulated genes according to RegulonDB (55) with expression higher than a threshold (10) across all replicates. Setting transcriptome quantification from the moderate lactose condition, we applied the adaptive-T prior shrinkage estimator “apeglm” and used Wald significance tests for detecting DEGs and obtaining the log_2_ fold changes (LFC2).

### 4.4. Metabolic pathway enrichment analysis

#### 4.4.1. Enrichment analysis by FGSEA

At the time of analysis, there were 368 pathways in EcoCyc for B REL606. 6 pathways that are not apparently regulated by gene products were discarded. Pathway enrichment was analyzed using FGSEA (58). Differentially expressed genes were pre-ranked by their log_2_ fold change. Pathway gene sets were defined with reference to the EcoCyc database. The pre-ranked gene data and pathway gene sets were then processed by the fgseaMultilevel function to obtain final results. Minimum gene set size was set to 3 with 200 bootstrap replicates. The threshold for pathway significance was 0.05.

Pathways in Figure 7 are linked by key metabolites and their flow direction. The pathway ontology in Figure 7 is annotated in the Ecocyc database. The pathway link and pathway ontology was extracted from EcoCyc and visualized with Cytoscape (59).

#### 4.4.2. Pathway regulatory mechanism similarity

To calculate the similarity between pathway regulation in different conditions, we used Spearman’s ρ rank correlation between the differential expression profile for each pathway. The similarity score for cases where all gene enrichment in a pathway are of the same direction is set to 1.

Metabolic pathway visualization used the Python package Escher (60).

### 4.5. Code availability

Analysis code is available at https://github.com/jcjray/ecoli_divergent_stress_pipeline

## Acknowledgments

We thank Jaden Anderson, Jimmy Budiardjo, Hung Do, Susan Egan, Chad Highfill, Erik Lundquist, Jacqueline J. Stevens, and Pinakin R. Sukthankar for valuable discussions and assistance. This project was supported by Institutional Development Awards (IDeA) from the National Institute of General Medical Sciences of the National Institutes of Health under grant numbers P20GM103418 and P20GM103638. The content is solely the responsibility of the authors and does not necessarily represent the official views of the National Institute of General Medical Sciences or the National Institutes of Health. Research reported in this publication was also made possible in part by the services of the KU Genome Sequencing Core. This core lab is supported by the National Institute of General Medical Sciences (NIGMS) of the National Institutes of Health under award number P20GM103638.

## Supplementary Figures

**Figure S1.**
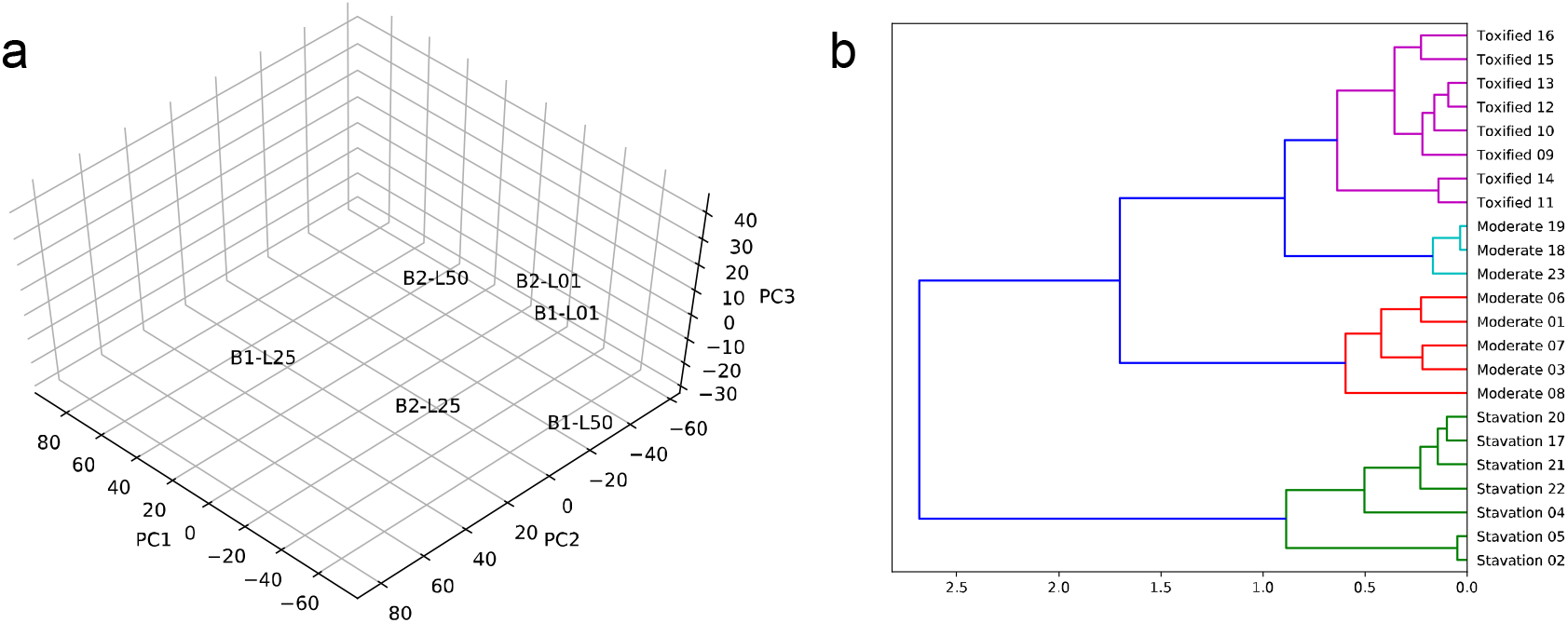
Data quality verification. **a**. PCA analysis for the transcriptome profile. **b**. hierarchical clustering using Wald significance. The clustering recapitulated treatment conditions.

**Figure S2.**
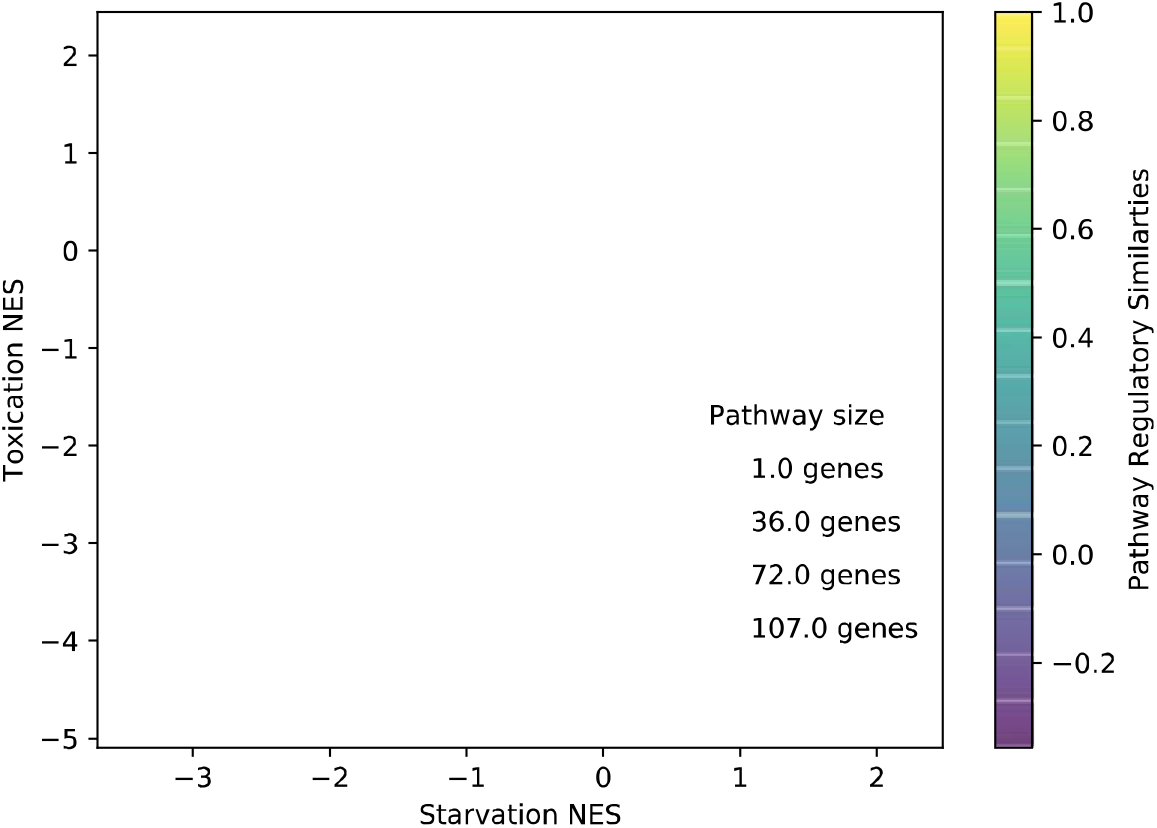
Pathway enrichment analysis overview. The plot is color-coded for the similarity of pathway regulation between starvation and toxicity. The size of the circle represents pathway gene set size, and the location of the bubbles are the normalized enrichment scores for each pathway.

**Figure S3.**
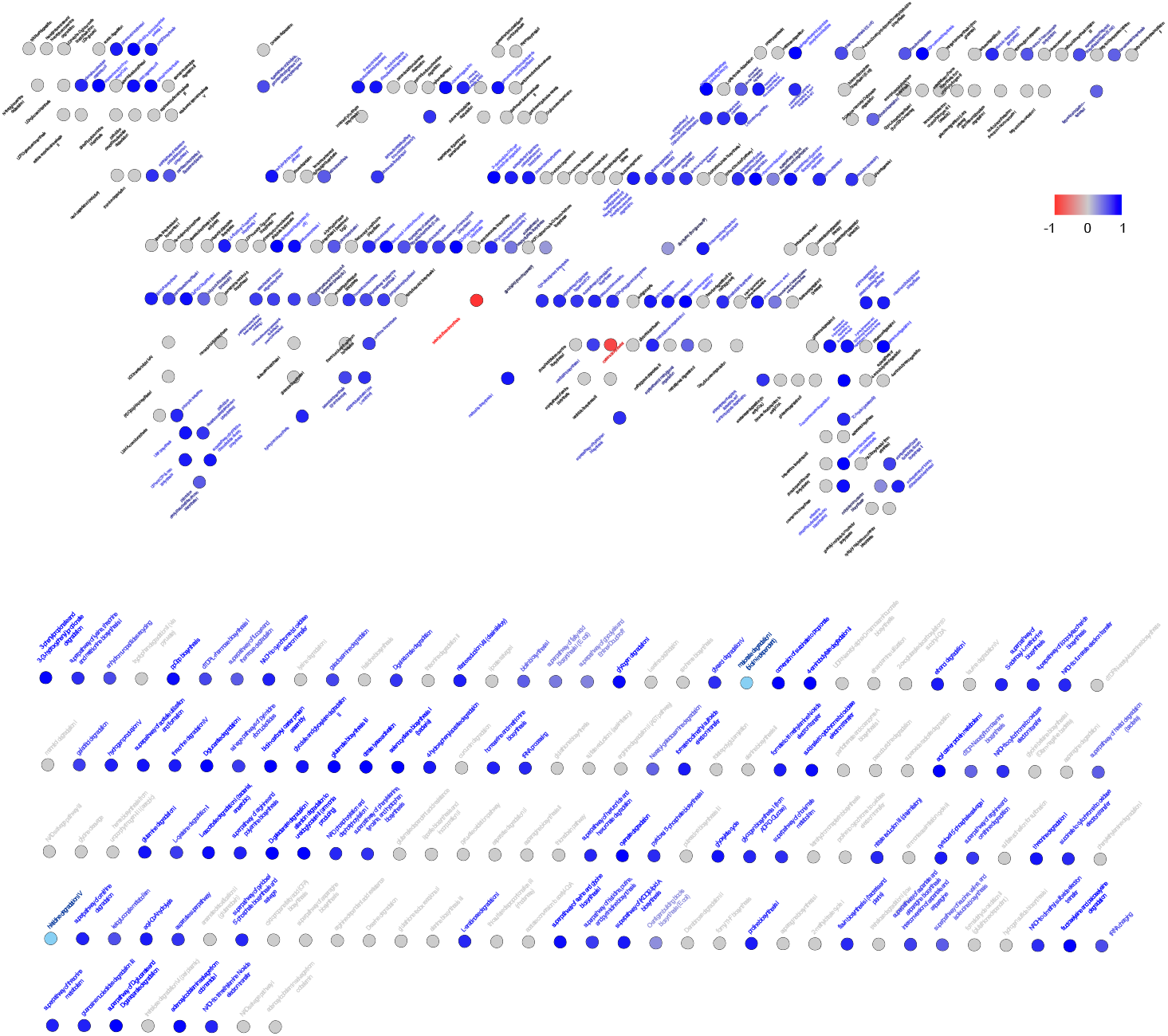
Pathway regulatory correlations between starving and toxified cells. Blue dots show a conserved regulatory regime for both stresses, and red dots show oppositely regulated pathways.

## Supplementary Tables

**Table S1.** Pathways enriched in both starvation and toxification. Pathway ID and name are according to EcoCyc database annotations, and normalized enrichment score is calculated using the R package fgsea.

| Pathway ID | Name | Gene Set Size | Spearman correlation | Toxication NES | Starvation NES |
| --- | --- | --- | --- | --- | --- |
| PUTDEG-PWY | putrescine degradation I | 2 | -1 | 1.48002492 | 4.65997042 |
| PWY0-1305 | glutamate dependent acid resistance | 2 | -1 | 1.1859878 | 1.59212915 |
| PWY0-461 | lysine degradation I | 2 | -1 | 1.35155215 | 1.83928035 |
| PWY0-1317 | L-lactaldehyde degradation (aerobic) | 2 | -1 | 1.46805444 | 2.01581953 |
| THRDLCTCAT-PWY | threonine degradation III (to methylglyoxal) | 2 | -1 | 1.46828561 | 1.77182093 |
| PWY0-1280 | ethylene glycol degradation | 2 | -1 | 2.32846128 | 2.44590459 |
| GLYCOLATEMET-PWY | glycolate and glyoxylate degradation I | 7 | 0.32142857 | 1.88244895 | 1.86338914 |
| GLYCOL-GLYOXDEG-PWY | superpathway of glycol metabolism and degradation | 11 | 0.36446564 | 1.88372208 | 2.02611816 |
| PWY0-42 | 2-methylcitrate cycle I | 6 | 0.37142857 | 1.69070836 | 2.03799995 |
| PWY-6961 | L-ascorbate degradation II (bacterial, aerobic) | 9 | 0.83333333 | 2.11098325 | 1.79204893 |
| PWY-6772 | hydrogen production V | 7 | 0.85714286 | 1.88508666 | 1.77736595 |
| PWY0-301 | L-ascorbate degradation I (bacterial, anaerobic) | 8 | 0.95238095 | 1.83854381 | 1.72700654 |
| LYXMET-PWY | L-lyxose degradation | 7 | 0.96428571 | 1.93769922 | 1.80519956 |
| THREONINE-DEG2-PWY | threonine degradation II | 2 | 1 | 1.25911262 | 1.90509234 |
| GLUCONSUPER-PWY | D-gluconate degradation | 2 | 1 | 1.702271 | 2.3647267 |
| PWY-66 | GDP-L-fucose biosynthesis I (from GDP-D-mannose) | 2 | 1 | 2.99269776 | 2.42917244 |
| PWY0-661 | PRPP biosynthesis II | 2 | 1 | 2.88193561 | 1.72370551 |
| PWY-6894 | thiamin diphosphate biosynthesis I (E. coli) | 2 | 1 | 2.70271621 | 2.35363775 |
| PWY-7181 | pyrimidine deoxyribonucleosides degradation | 2 | 1 | 1.15009838 | 1.93460402 |
| PWY0-1300 | 2-O-alpha-mannosyl-D-glycerate degradation | 2 | 1 | 2.45899684 | 2.4412234 |
| 2PHENDEG-PWY | phenylethylamine degradation I | 2 | 1 | 1.97359284 | 2.09844355 |
| PWY0-1309 | chitobiose degradation | 2 | 1 | 1.88736692 | 1.19095321 |
| TRESYN-PWY | trehalose biosynthesis I | 2 | 1 | 1.83677266 | 3.00331018 |
| PWY-7247 | beta-D-glucuronide and D-glucuronate degradation | 2 | 1 | 1.82541516 | 1.60708482 |
| PWY-46 | putrescine biosynthesis III | 2 | 1 | 1.78260691 | 1.40424707 |
| PWY-6019 | pseudouridine degradation | 2 | 1 | 3.23424124 | 2.69738572 |
| XYLCAT-PWY | xylose degradation I | 2 | 1 | 2.34239671 | 3.07625823 |
| PWY0-1466 | trehalose degradation VI (periplasmic) | 2 | 1 | 1.70019343 | 4.27176029 |
| PWY0-1477 | ethanolamine utilization | 2 | 1 | 2.32607631 | 1.9938891 |
| GLUCARGALACTSUPER-PWY | superpathway of D-glucarate and D-galactarate degradation | 5 | 1 | 1.85197625 | 1.98616611 |
| AST-PWY | arginine degradation II (AST pathway) | 5 | 1 | 1.67181348 | 2.15994831 |
| GLUCARDEG-PWY | D-glucarate degradation I | 4 | 1 | 1.69740595 | 1.84433906 |
| GALACTARDEG-PWY | D-galactarate degradation I | 4 | 1 | 1.69740595 | 1.89776655 |
| PWY0-541 | cyclopropane fatty acid (CFA) biosynthesis | 1 | NC | 1.49596218 | 1.94296286 |
| TREDEGLOW-PWY | trehalose degradation I (low osmolarity) | 1 | NC | 2.18675062 | 1.41455366 |
| PWY0-1315 | L-lactaldehyde degradation (anaerobic) | 1 | NC | 2.32846128 | 1.49622475 |
| PWY-6476 | cytidyl molybdenum cofactor biosynthesis | 1 | NC | 2.73758113 | 6.40687046 |
| TRYPDEG-PWY | tryptophan degradation II (via pyruvate) | 1 | NC | 2.71945032 | 2.32442325 |
| PWY0-1306 | L-galactonate degradation | 1 | NC | 2.65064294 | 2.07270313 |
| PWY0-1313 | acetate conversion to acetyl-CoA | 1 | NC | 1.19407481 | 4.21849854 |
| PYRUVOX-PWY | pyruvate oxidation pathway | 1 | NC | 1.1743514 | 3.66245495 |
| PWY0-1527 | curcumin degradation | 1 | NC | 2.09016159 | 3.65971023 |
| PWY-4621 | arsenate detoxification II (glutaredoxin) | 1 | NC | 1.39610105 | 1.14732764 |
| SORBDEG-PWY | D-sorbitol degradation II | 1 | NC | 1.25727254 | 1.08246131 |

**Table S2.**
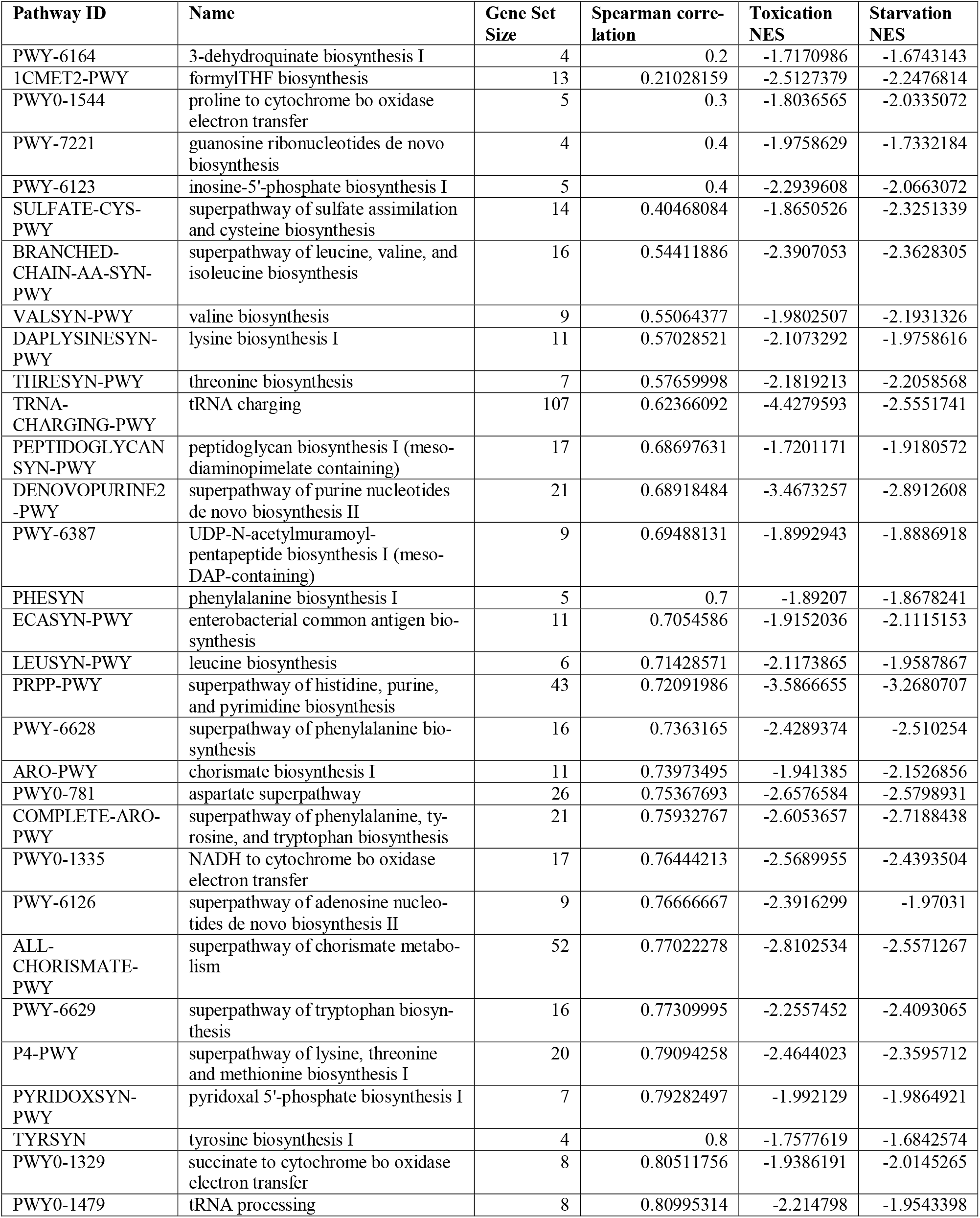
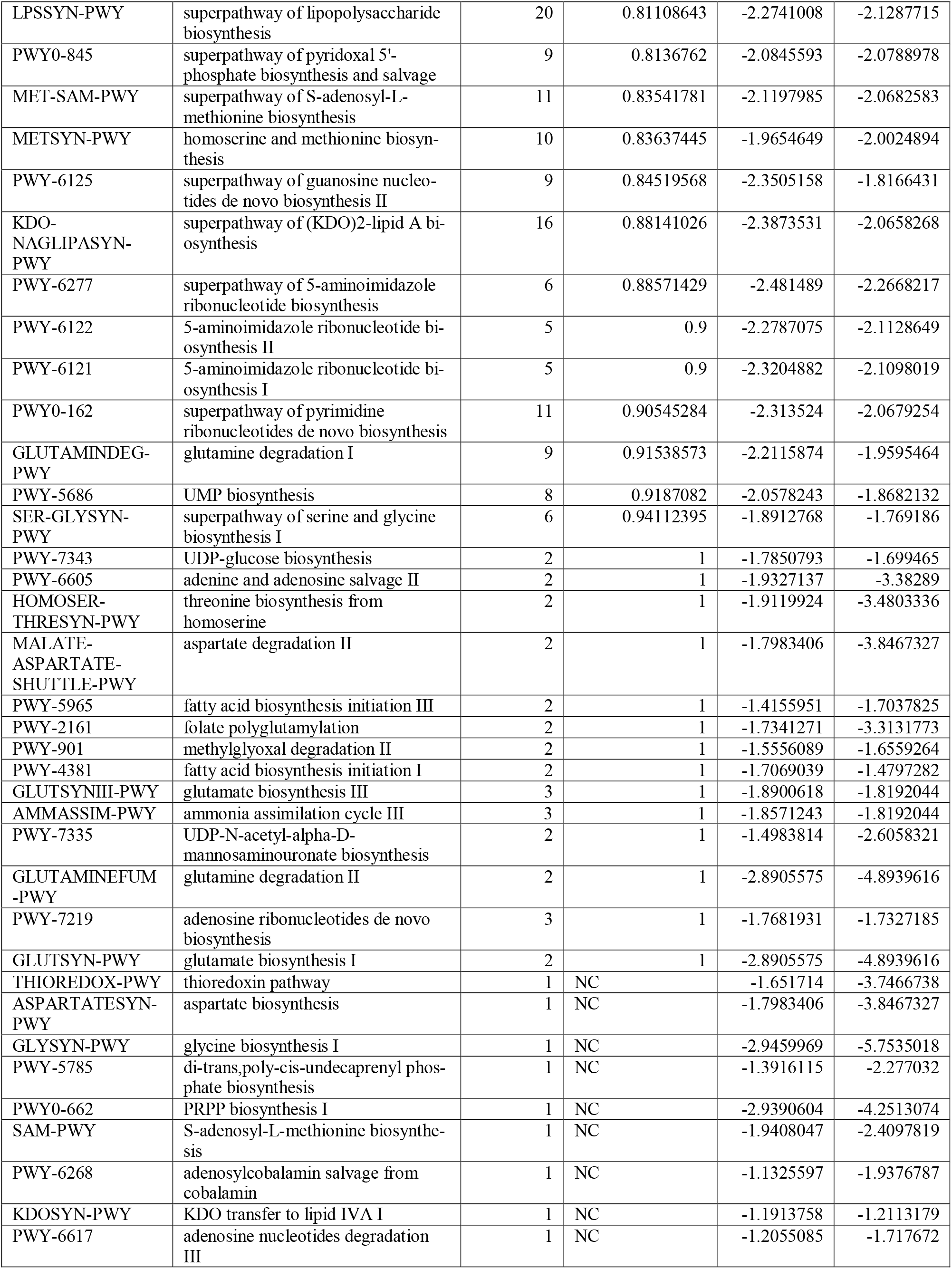
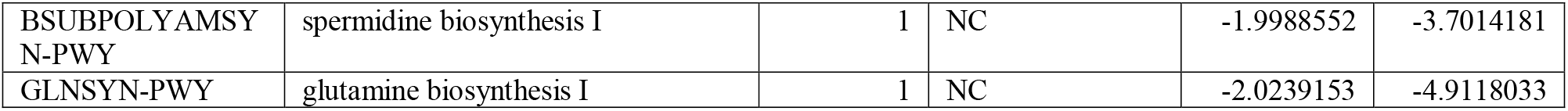
Pathways that are downregulated in both starvation and toxification. Pathway ID and name are according to EcoCyc database annotations, and normalized enrichment score is calculated using the R package fgsea.

**Table S3.** Pathways uniquely enriched in starvation. Pathway ID and name are according to EcoCyc database annotations, and normalized enrichment score is calculated using the R package fgsea.

| Pathway ID | Name | Gene Set Size | Spearman correlation | Toxication NES | Starvation NES |
| --- | --- | --- | --- | --- | --- |
| FAO-PWY | fatty acid beta-oxidation I | 9 | -0.0666667 | 0 | 2.22387757 |
| PWY-4261 | glycerol degradation I | 5 | 0.5 | 0 | 2.03254717 |
| PWY0-1182 | trehalose degradation II (trehalase) | 3 | 0.5 | 0 | 1.62706335 |
| PWY-6952 | glycerophosphodiester degradation | 6 | 0.71428571 | 0 | 1.79463742 |
| PWY0-381 | glycerol and glycerophosphodiester degradation | 7 | 0.75678747 | 0 | 1.92648052 |
| GLYOXDEG-PWY | glycolate and glyoxylate degradation II | 5 | 0.9 | 0 | 1.93846673 |
| PWY0-1299 | arginine dependent acid resistance | 2 | 1 | 0 | 1.19479501 |
| DARABCAT-PWY | D-arabinose degradation II | 1 | NC | 0 | 1.26124042 |
| PWY-7346 | UDP-alpha-D-glucuronate biosynthesis (from UDP-glucose) | 1 | NC | 0 | 2.0622656 |
| PWY0-1337 | oleate beta-oxidation | 1 | NC | 0 | 3.74727095 |
| MANNIDEG-PWY | mannitol degradation I | 1 | NC | 0 | 3.20579409 |
| GLUAMCAT-PWY | N-acetylglucosamine degradation I | 2 | NC | 0 | 1.21830745 |
| PWY-7179 | purine deoxyribonucleosides degradation | 2 | NC | 0 | 1.39473893 |
| PWY-6028 | acetoin degradation | 2 | NC | 0 | 1.47252358 |

**Table S4.**
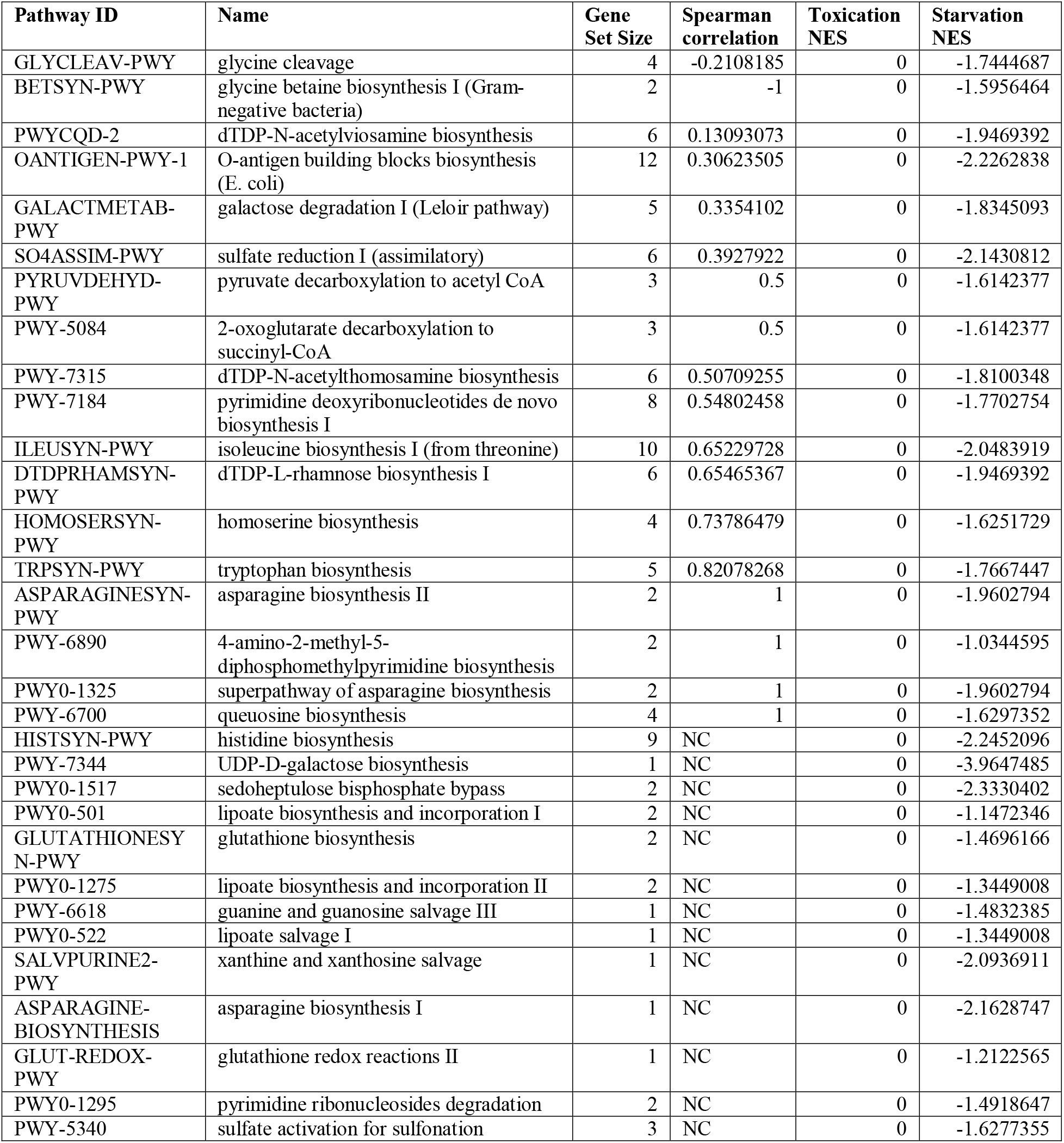
Pathways that are uniquely downregulated in starvation. Pathway ID and name are according to EcoCyc database annotations, and normalized enrichment score is calculated using the R package fgsea.

**Table S5.** Pathways that are uniquely enriched in toxicity. Pathway ID and name are according to EcoCyc database annotations, and normalized enrichment score is calculated using the R package fgsea.

| Pathway ID | Name | Gene Set Size | Spearman correlation | Toxicity NES | Starvation NES |
| --- | --- | --- | --- | --- | --- |
| PWYCQD-3 | N-acetyl-galactosamine degradation | 9 | 0.51666667 | 1.84151128 | 0 |
| PWYCQD-4 | galactosamine degradation | 8 | 0.66666667 | 1.9179961 | 0 |
| GALACTITOLCAT-PWY | galactitol degradation | 5 | 0.7 | 1.59677363 | 0 |
| PWY0-1321 | nitrate reduction III (dissimilatory) | 12 | 0.8041958 | 1.90376903 | 0 |
| 3-HYDROXYPHENYLACETATE-DEGRADATION-PWY | 4-hydroxyphenylacetate degradation | 8 | 0.88095238 | 1.73575101 | 0 |
| PWY0-1356 | formate to dimethyl sulfoxide electron transfer | 9 | 0.88333333 | 1.69948976 | 0 |
| PWY-6690 | cinnamate and 3-hydroxycinnamate degradation to 2-oxopent-4-enoate | 8 | 0.92857143 | 1.72900525 | 0 |
| HCAMHPDEG-PWY | 3-phenylpropanoate and 3-(3-hydroxyphenyl)propanoate degradation to 2-oxopent-4-enoate | 8 | 0.92857143 | 1.72900525 | 0 |
| PWY0-1277 | 3-phenylpropionate and 3-(3-hydroxyphenyl)propionate degradation | 12 | 0.97022916 | 1.92679151 | 0 |
| RHAMCAT-PWY | L-rhamnose degradation I | 4 | 1 | 1.58032918 | 0 |
| PWY0-1465 | D-malate degradation | 1 | NC | 1.36929171 | 0 |
| PWY0-981 | taurine degradation IV | 1 | NC | 1.33115815 | 0 |

**Table S6.**
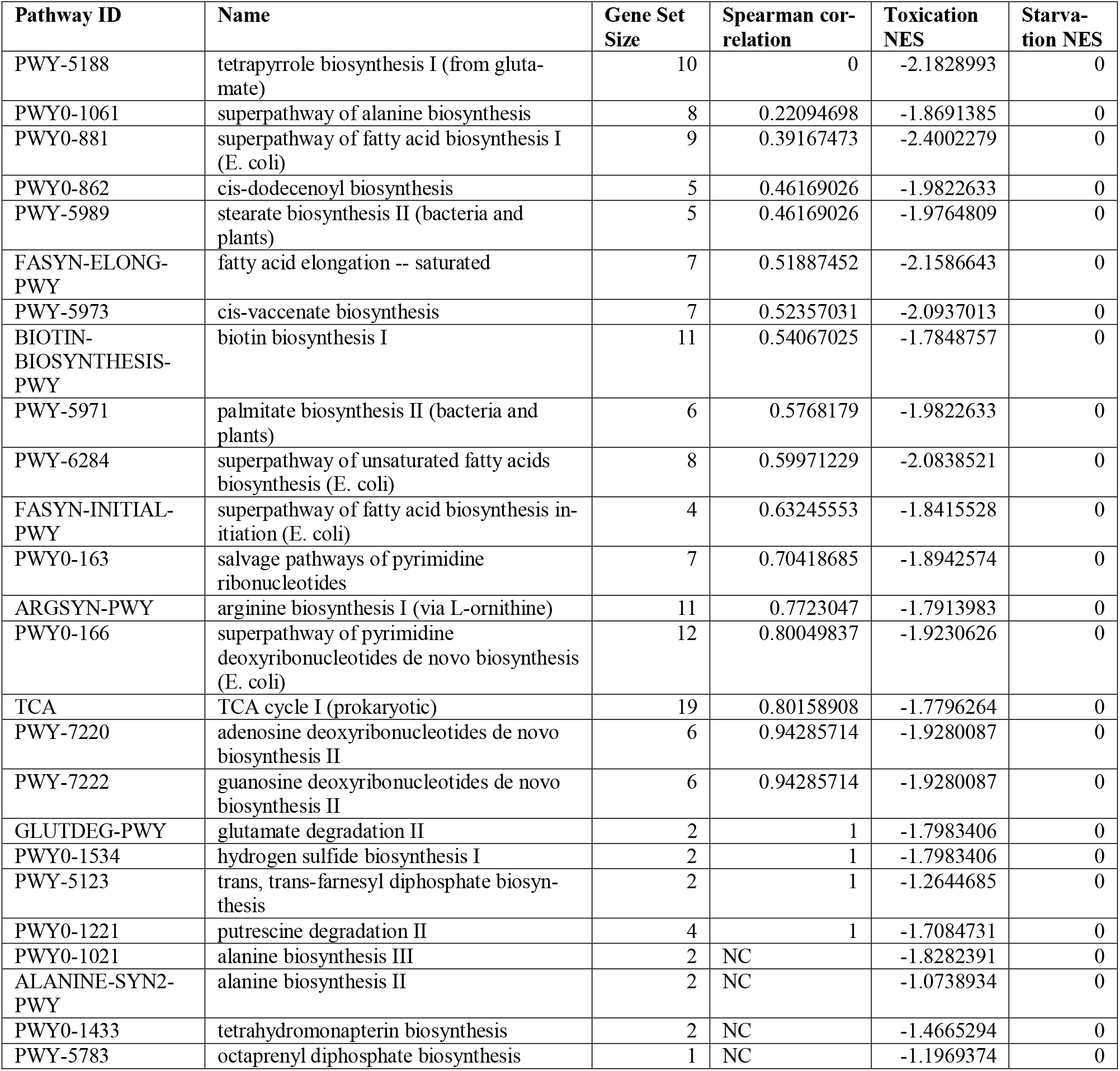
Pathways that are uniquely downregulated in toxification condition. Pathway ID and name are consistent with EcoCyc database annotations, and normalized enrichment score is calculated with R package fgsea.

## Footnotes

1 Practical constraints prevented our study from creating a distance metric for how quantitative changes in lactose correspond to quantitative changes in gene expression beyond the cases shown here. Interpretations are thus carefully limited to categorical comparisons between the culture states rather than quantitative shifts of gene expression from changes in lactose concentration.

